# Neutral Drift and Threshold Selection Promote Phenotypic Variation

**DOI:** 10.1101/2023.04.05.535609

**Authors:** Ayşe N. Erdoğan, Pouria Dasmeh, Raymond D. Socha, John Z. Chen, Ben Life, Rachel Jun, Linda Kiritchkov, Dan Kehila, Adrian W.R. Serohijos, Nobuhiko Tokuriki

## Abstract

Phenotypic variations within a population exist on different scales of biological organization and play a central role in evolution by providing adaptive capacity at the population-level. Thus, the question of how evolution generates phenotypic variation within an evolving population is fundamental in evolutionary biology. Here we address this question by performing experimental evolution of an antibiotic resistance gene, VIM-2 β-lactamase, combined with diverse biochemical assays and population genetics. We found that neutral drift, *i.e.*, evolution under a static environment, with a low antibiotic concentration can promote and maintain significant phenotypic variation within the population with >100-fold differences in resistance strength. We developed a model based on the phenotype-environment-fitness landscape generated with >5,000 VIM-2 variants, and demonstrated that the combination of “mutation-selection balance” and “threshold-like fitness-phenotype relationship” is sufficient to explain the generation of large phenotypic variation within the evolving population. Importantly, high-resistance conferring variants can emerge during neutral drift, without being a product of adaptation. Our findings provide a novel and simple mechanistic explanation for why most genes in nature, and by extension, systems and organisms, inherently exhibit phenotypic variation, and thus, population-level evolvability.

## Introduction

A population often encounters and needs to adapt to environmental changes; otherwise, it could face extinction. Although the influx of new mutations can provide inheritable phenotypic change, they may be insufficient when environmental perturbations are sudden and large, therefore necessitating the pre-existence of phenotypic variation in natural populations^1,2^. Indeed, phenotypic variation at different scales of biological organizations, including at the protein level, is commonly observed both within populations and within species as a whole, pointing towards the importance of existence of such diversity as a major mechanism for providing evolving populations adaptive capacity, *i.e.,* population-level evolvability^1,3–6^. For example, the pre-existence of gene variants that confer different levels of antibiotic resistance in bacteria allows for the survival of the bacterial population upon a sudden increase in antibiotic concentration, where bacteria with high-resistance variants can rescue the population^7^. However, even though the importance of phenotypic variation is indeed well recognized, our understanding of the mechanisms for its generation and maintenance is limited^8^.

Phenotypic variation is often regarded as a by-product of “differential selection”, due to spatial and/or temporal environmental fluctuations^8^. It is also known that phenotypic traits that are not directly under selection pressure tend to exhibit variation, so-called hidden phenotypic variation^5,8–12^. However, these theories apply to only specific and limited evolutionary scenarios, and they cannot provide an explanation for the ubiquity of phenotypic variation observed in nature. For instance, recent experimental evolution studies demonstrated that bacterial populations evolved under constant, low-level antibiotic exposure resulting in heterogenetic antibiotic resistance phenotypes, with a fraction of variants conferring high levels of resistance^13–16^. While these studies suggest that evolution in a simple and static environment may lead to phenotypic variation, the underlying mechanism for this is yet unknown.

Protein evolution under purifying selection in a static environment can be referred as “neutral drift”, where the accumulation of mutations occurs while the functionality of the protein is sustained above a certain threshold to maintain organismal fitness^17–20^. It is known that neutral drift promotes genotypic variation in proteins^17,18^. Nonetheless, these mutations can be non-neutral in their effect on the protein phenotype. This can also result in variation in non-physiological and promiscuous functions of the protein as they are not directly under the selection pressure^20–24^. However, it is unclear if neutral drift can generate and maintain phenotypic variation for the trait that is directly under selection pressure. If so, this provides a robust mechanistic explanation for the phenotypic variations that are commonly observed within a population and species, as most proteins have undergone neutral drift.

In this work, we addressed the question by conducting an evolutionary experiment on VIM-2 β-lactamase. We evolved VIM-2 under different evolutionary scenarios including neutral drift (**Fig. 1**). We tracked changes in genotypic and phenotypic diversity within the population and examined evolutionary conditions that promote high phenotypic variation. Then, using an empirically obtained fitness landscape for VIM-2^25,26^, we simulated the evolution of VIM-2 and demonstrated that mutation-selection balance combined with a simple threshold relationship between the phenotype and fitness can easily result in the generation and maintenance of phenotypic diversity within an evolving population.

**Fig. 1.**
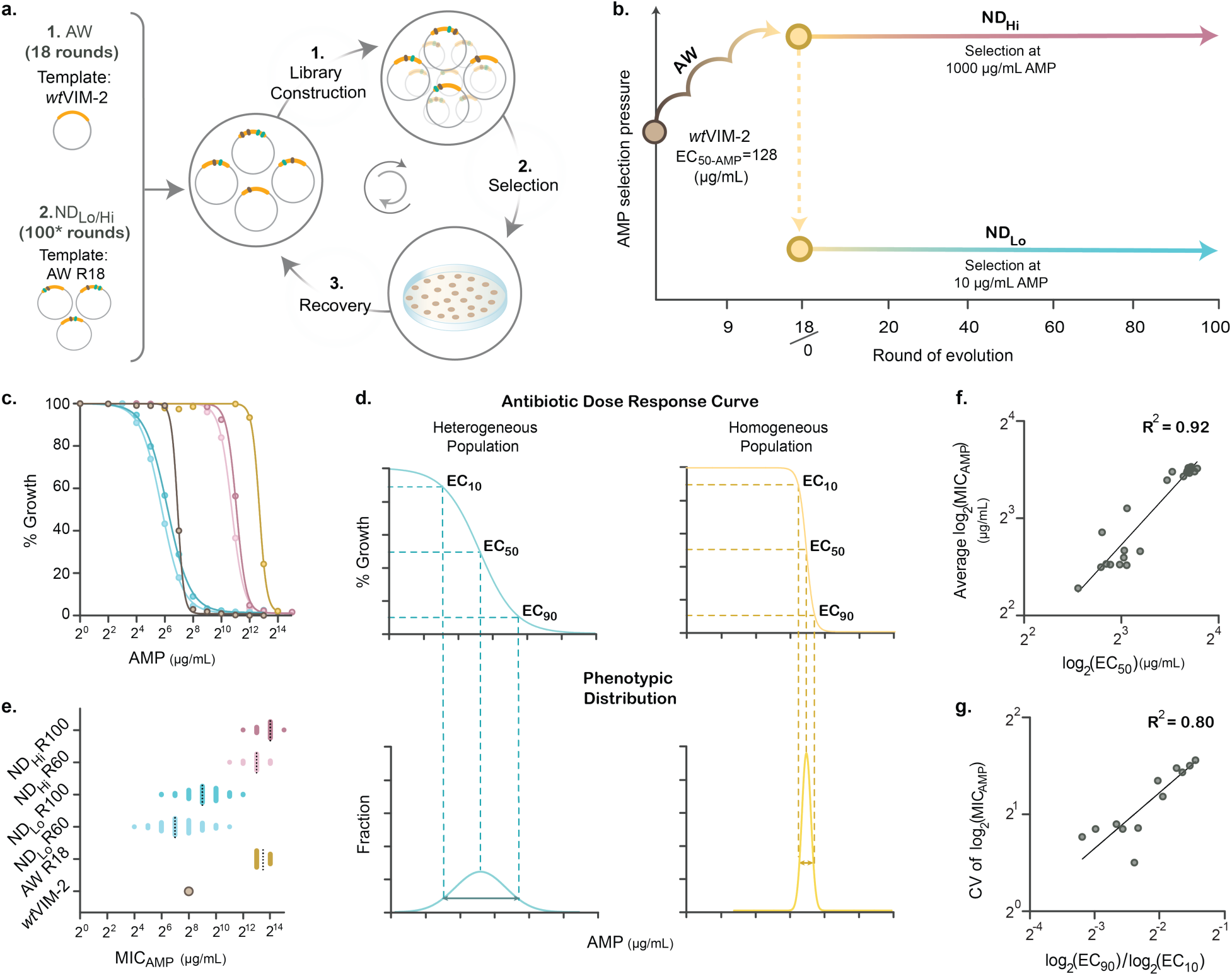
Scheme of experimental evolution of VIM-2 and analysis pipeline. **a**, Overview of experimental evolution. **b,** Scheme of the ampicillin concentrations used in the experimental evolution: During adaptive walk (AW), ampicillin concentration in the selection media is gradually increased. After AW, the population is split into two lineages and then subjected to 100 iterative rounds of neutral drift with high (1,000 µg/mL ampicillin, NDHi) and low (10 µg/mL ampicillin, NDLo) concentrations of ampicillin. **c**, Representative population-level phenotypes (dose-response curves) Refer to panel **e** for the colouring scheme. **d**, Illustration of how dose-response curves reflect the phenotypic variation in the population. **e**, Representative phenotype measurements (MIC) of individual variants sampled along each trajectory. **f**, **g**, Correlation between characteristics from the population-level phenotypic assay (dose-response curve), and individual-level assays (MIC): MICAMP values *versus* EC50 (**f**) and coefficient of variation (CV) of log2-transformed MICAMP values *versus* log2-transformed EC90/EC10 (**g**).

## Results

### Experimental evolution of VIM-2 β-lactamase and characterizations of the populations

We conducted experimental evolution of VIM-2 by expressing it in an *Escherichia coli* strain (E. cloni 10G), and applying selection on agar plates with the required ampicillin concentration (**Fig. 1a)**. The minimum inhibitory concentration on an agar plate (MIC) of the *E. coli* strain without expressing any VIM-2 variant was 4 µg/mL ampicillin, while expressing wild-type VIM-2 (*wt*VIM-2) conferred 32-fold higher resistance to the strain (MIC=128 µg/mL). *wt*VIM-2 was first subjected to 18 rounds of directed evolution for higher antibiotic resistance (adaptive walk or AW) by gradually increasing the ampicillin concentration until the ampicillin resistance of the population was plateaued (at 4,096 µg/mL, **Fig. 1b** and **Supp. Data 1**). This concentration was an apparent limit for the evolution since selection with the next 2-fold increase in concentration, 8,192 µg/mL ampicillin, did not yield any colonies after R12. Subsequently, the adapted population of VIM-2 variants (*i.e.,* AW-R18) was divided into two lineages and further subjected to 100 rounds of neutral drift (ND) under a static environment (**Fig. 1b**). One linage, ND_Hi_, was evolved with a high ampicillin concentration (1,000 µg/mL). The other linage, ND_Lo_, was subjected to a low ampicillin concentration (10 µg/mL). In this way, we examined the behaviour of the same starting population under different evolution scenarios.

The phenotype of a population was determined by antibiotic dose-response growth assays on the *E. coli* population expressing each VIM-2 library (**Fig. 1c)**. The dose-response curve was fitted to a sigmoidal function to obtain the effective concentrations of antibiotic that inhibits the growth of 10, 50 and 90% (EC_10_, EC_50_, EC_90_) of the population (**Fig. 1d** and **Extended Data Fig. 1**). Each library contained over 10,000 VIM-2 variants, in which the EC_50_ provided an estimate for the median resistance of the population. Meanwhile, EC_90_/EC_10_, the fold difference between the upper and lower bound of the curve’s transition range, reflected the variation in the distribution of resistance phenotypes within the population (**Fig. 1e**); EC_90_/EC_10_ for a monoclonal population was typically <2.5 (**Fig. 1d**, **Supp. Table 1**). Also, the MIC of representative VIM-2 variants was determined to monitor variation within each population. Importantly, the dose-response curve recapitulated the phenotypic characteristics of the populations, as EC_50_ and EC_90_/EC_10_ were highly correlated to the mean and variation in MICs of isolated variants, respectively (**Fig. 1f-g**).

### Neutral drift at a low antibiotic selection threshold promoted and maintained phenotypic variation

We examined the evolving populations along each trajectory (AW, ND_Hi_, ND_Lo_) for three key parameters: ***i*)** the genetic variation, ***ii*)** the apparent selection pressure, and ***iii*)** the resulting phenotypic variation (**Fig. 2**). Over the course of the evolution, mutations were consistently accumulated, and genetic variation within the populations continuously increased, in particular during neutral drift (**Fig 2a**, and **Extended Data Fig. 2**). Genetic diversity of ND_Lo_ (∼25% divergence from *wt*VIM-2 at R100) was only moderately higher than that of ND_Hi_ (∼20% at R100). We estimated the selection pressure using the ratio between the amino acid (*N*_a_) and nucleotide (*N*_t_) mutations, where *N*_a_/*N*_t_ = 0.46 for a random walk accepted all nucleotide mutations (**Methods**). During AW, the *N*_a_/*N*_t_ ratio reached as much as 0.54 in the early rounds, higher than the random walk ratio, suggesting the enrichment of adaptive and nonsynonymous mutations over synonymous mutations for higher ampicillin resistance (**Fig. 2b**). Meanwhile, throughout most of ND_Hi/Lo_, *N*_a_/*N*_t_ exhibited a decreasing trend, dropping and remaining below 0.46 for most of ND_Hi_ and the latter half of ND_Lo_. This indicated that both populations underwent purifying selection, whereby nonsynonymous mutations with negative effects were mostly purged to maintain the resistance phenotypes of the variants above the selection threshold (**Fig. 2b**). Nonetheless, ND_Hi_ exhibited consistently lower *N*_a_/*N*_t_, throughout the evolution compared to that of ND_Lo_, which confirms slightly higher selection stringency in the ND_Hi_ population.

**Fig. 2.**
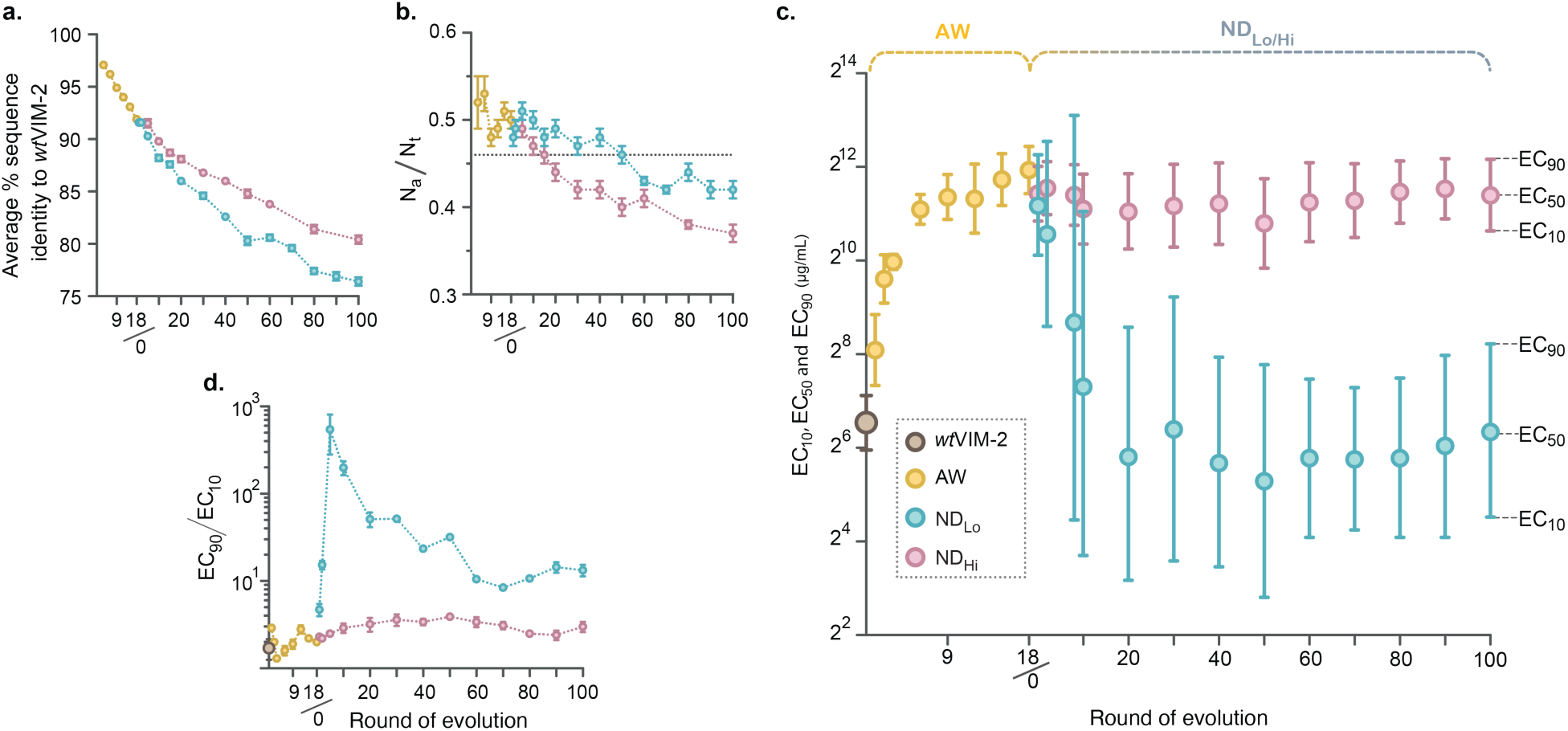
Phenotypic and genotypic characteristics of the AW, NDLo and NDHi libraries. **a**, Changes in amino acid sequence identity shared with *wt*VIM-2. **b**, Na/Nt ratios throughout the evolving trajectories. The Na/Nt of a random walk (0.46) is denoted by the horizontal line (**Methods**). **c**, Changes in EC10, EC50 and EC90 of the evolved VIM-2 libraries. The central dot represents EC50, and bars at both ends for EC10 and EC90 respectively. **d**, Changes in EC90/EC10 during the evolution, error bars represent standard error between 4 biological replicates for the NDLo populations and 2 biological replicates for the AW and NDH populations. The data used for panel **a** and panel **b** are shown in **Supplementary Table 2;** the data used for panels **c** and **d** are shown in **Supplementary Table 3.**

Despite the similarity observed at the genetic level, the phenotypic variation was substantially different between trajectories (**Fig. 2c, d**). During AW, EC_50_ gradually increased by 40-fold (**Fig. 2c**), but the phenotypic variation, gauged by the magnitude of EC_90_/EC_10_, was consistently as low as a monoclonal population, homogenous with respect to resistance levels (**Fig. 1d, Fig. 2d)**. In agreement with this, individual variants picked randomly from the AW populations exhibited similar resistance levels (**Fig. 2d** and **Extended Data Fig. 1**). During the subsequent 100 rounds of neutral drift, ND_Hi_ maintained its high resistance trait (EC_50_ ∼2,500 µg/mL) and narrow phenotypic variation (EC_90_/EC_10_ <4), due to the selection against all variants that do not confer high levels of resistance (*i.e.,* >1,000 ug/mL) (**Fig. 2c-d** and **Extended Data Fig. 1**). On the contrary, the ND_Lo_ population exhibited a more dynamic trajectory. As expected, EC_50_ gradually decreased from ∼4,000 µg/mL in the first 20 rounds to just above the purifying selection threshold (∼50 µg/mL). Then, EC_50_ remained at the same level in the next 80 rounds (**Fig. 2c**). Phenotypic variation of ND_Lo_ radically increased in the first 10 rounds (EC_90_/EC_10_ >500 in R10), with the mixture of parental high-resistance variants and emerging medium and low resistance variants (**Fig. 2d** and **Extended Data Fig. 1**). This was anticipated as ND_Lo_ would initially allow the accumulation of mildly negative mutations that decrease resistance, which reflects “relaxed purifying selection”^27,28^. Intriguingly, after R20, when EC_50_ became constant, EC_90_/EC_10_ ratio only moderately decreased and remained high at ∼10 in the next 80 rounds (**Fig 2c**). The characterization of individual variants also showed large MIC variations in the ND_Lo_ populations with a >200-fold difference in MIC, and some variants conferring >100-fold higher MICs than the selection threshold of 10 µg/mL (**Fig. 1c** and **Extended Data Fig. 1**). These observations suggest that high variation of antibiotic resistance levels can still be maintained in evolving populations subjected to neutral drift at a low antibiotic selection threshold.

### Enhanced phenotypic variation for non-selected antibiotics

We further investigated how the genotypic variation affected phenotypes that were not directly under selection. We measured the resistance conferred by variants along each trajectory to two additional non-selected classes of β-lactam antibiotics: cefotaxime and meropenem (**Fig. 3** and **Extended Data Fig. 3**). The ND_Hi_ populations showed significantly higher variation in the resistance levels against (∼4 fold) both cefotaxime and meropenem compared to ampicillin (**Extended Data Fig. 4g**). The phenotypic variation observed for resistance for non-selected antibiotics in the ND_Lo_ populations was similar to the variation for ampicillin in ND_Lo_ but much higher than variation in ND_Hi_ (**Extended Data Fig. 4h**). Interestingly at the level of individual variants, the cefotaxime and meropenem resistance deviated from that of ampicillin and between each other to some extent (**Fig. 3a-d,** and **Extended Data Fig. 4a-f**). This uncoupling of mutational effects on three antibiotics generated further phenotypic variation within the populations, *e.g.*, the substrate specificity between ampicillin and cefotaxime resistance varies >1,000-fold among variants within the same population (**Fig. 3d**). Thus, neutral drift expands phenotypic diversity and evolvability of the antibiotic resistance gene population beyond the antibiotics used for selection.

**Fig. 3.**
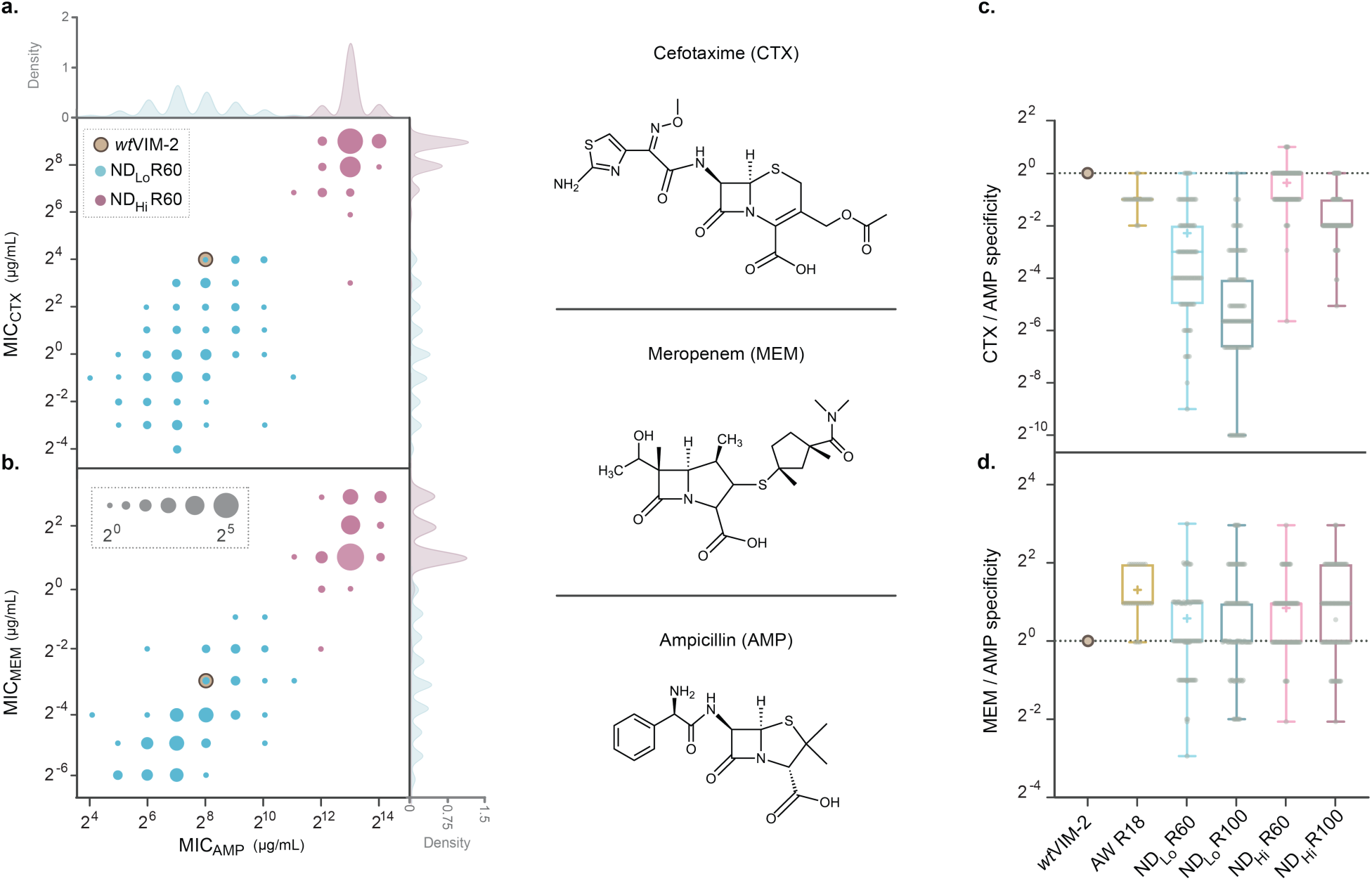
The distribution of resistance against selected and non-selected antibiotics. Fig. 3. The distribution of resistance against selected and non-selected antibiotics. **a, b.** The correlation between resistances between selected antibiotics (ampicillin, MICAMP) and non-selected antibiotics (cefotaxime, MICCTX and meropenem, MICMEM) of 92 variants from the R60 NDLo and NDHi populations. The resistance level was measured as MIC in the agar plate assays. The radii of the circles are weighted by the number of variants with the same substrate specificity. The Gaussian kernel density distribution of variants that have a certain AMP, CTX and MEM MIC is shown on the top and right axes in lighter grey (bandwidth=0.02). *wt*VIM-2’s resistance profile was indicated with a tan circle with dark brown edges. Chemical structures of the β-lactam antibiotics used in characterizing the resistance phenotype of VIM-2 variants is given to the right. **c, d**. The distribution of the relative substrate specificity, the resistance level against non-selected antibiotics: CTX (**c**), and MEM (**d**) over selected antibiotics, AMP, normalized over the same ratio for *wt*VIM-2 (shown as dashed lines, refer to **Methods** for the formula). The correlation between MICAMP, MICCTX, and MICMEM for the R60 populations and the comparison of fold-differences between MICAMP, MICCTX, and MICMEM for the evolved enzyme populations are shown in **Extended Data Fig. 4**.

### Phenotypic variation in ND_Lo_ was not caused by sub-MIC selection or evolutionary hysteresis

We then investigated the mechanism underlying the emergence and maintenance of the observed phenotypic variation in the ND_Lo_ populations. To this end, we hypothesized several potential mechanisms, and examined their likelihood. The first possibility was that a low ampicillin concentration provides some selective advantage to high-resistance variants over lower-resistance variants *i.e*., sub-MIC selection^29,30^. However, our previous work on deep mutational scanning (DMS) of *wt*VIM-2 indicated otherwise^25,26^. The relationship between EC_50_ and fitness of >5,000 VIM variants can be described as a single sigmoid function with a steep slope, with no apparent advantage for high-resistant variants above the selection threshold^25,26^. Supporting this, when ND_Lo_ R60 and R100 populations were subjected to five cycles of selection under 10 µg/mL ampicillin (P10) without mutagenesis, the phenotypic variation remained the same (**Extended Data Fig. 5a**), confirming that sub-MIC selection was not critical for the phenotypic variation observed in the ND_Lo_ populations (**Extended Data Fig. 5b-c**). We also tested if there was a fitness cost for high-resistance variants, via an extended transformation experiment in the absence of ampicillin (P0) (**Extended Data Fig. 5a**). The phenotypic variation was also unchanged over five rounds of transformations, confirming that there was no significant difference in the fitness cost among the variants within the selected populations (**Extended Data Fig. 5d**).

The second hypothesis we tested was “evolutionary hysteresis”, whereby the historical changes in the selection pressure that the VIM-2 populations experienced might cause high-resistance variants to remain in the populations. Specifically, *wt*VIM-2 was initially evolved to its maximum resistance via AW, and subsequently subjected to neutral drift, and thus, some variants could have retained high resistance by accumulating only neutral mutations. However, the phenotypic variation was maintained over 80 rounds of neutral drift (over 60 amino acid mutations per variant on average). Thus, it is unlikely for variants to exclusively obtain neutral mutations and retain high resistance for so long when the selection pressure does not require it. Also, the phylogenetic analysis of variants showed scattered lineages of high-resistance variants across the ND_Lo_ phylogenetic tree as opposed to clustering of high resistance variants, suggesting that high-resistance variants emerged from low-resistance subpopulations during the neutral drift rather than being carried forward from high resistance ancestral variants (**Extended Data Fig. 6**).

Furthermore, to examine whether neutral drift is truly sufficient to promote phenotypic variation, we performed an additional line of ND experiment with 10 µg/mL ampicillin starting from *wt*VIM-2 (ND_Lo-wt_) (**Fig. 4a**). Importantly, *wt*VIM-2 exhibits an EC_50_ (100 µg/mL); 10-fold above the concentration used for the selection, and confers complete resistance to *E. coli* at 10 µg/mL. After only several rounds of neutral drift, ND_Lo-wt_ exhibited comparably high phenotypic variation to the ND_Lo_ populations (**Fig. 4a-c**). Peculiarly, ND_Lo-wt_ contained higher resistance variants (EC_50_>500 µg/mL) than *wt*VIM-2, suggesting high-resistance variants emerged during neutral drift without an adaptive selective force for high antibiotic concentrations **(Supplementary Table 6**). These observations confirmed that evolutionary hysteresis is not the cause for high phenotypic variation in ND_Lo_. On the contrary, these suggest that neutral drift with a low antibiotic concentration is sufficient to both maintain and generate high phenotypic variation, including higher resistance variants.

**Fig. 4.**
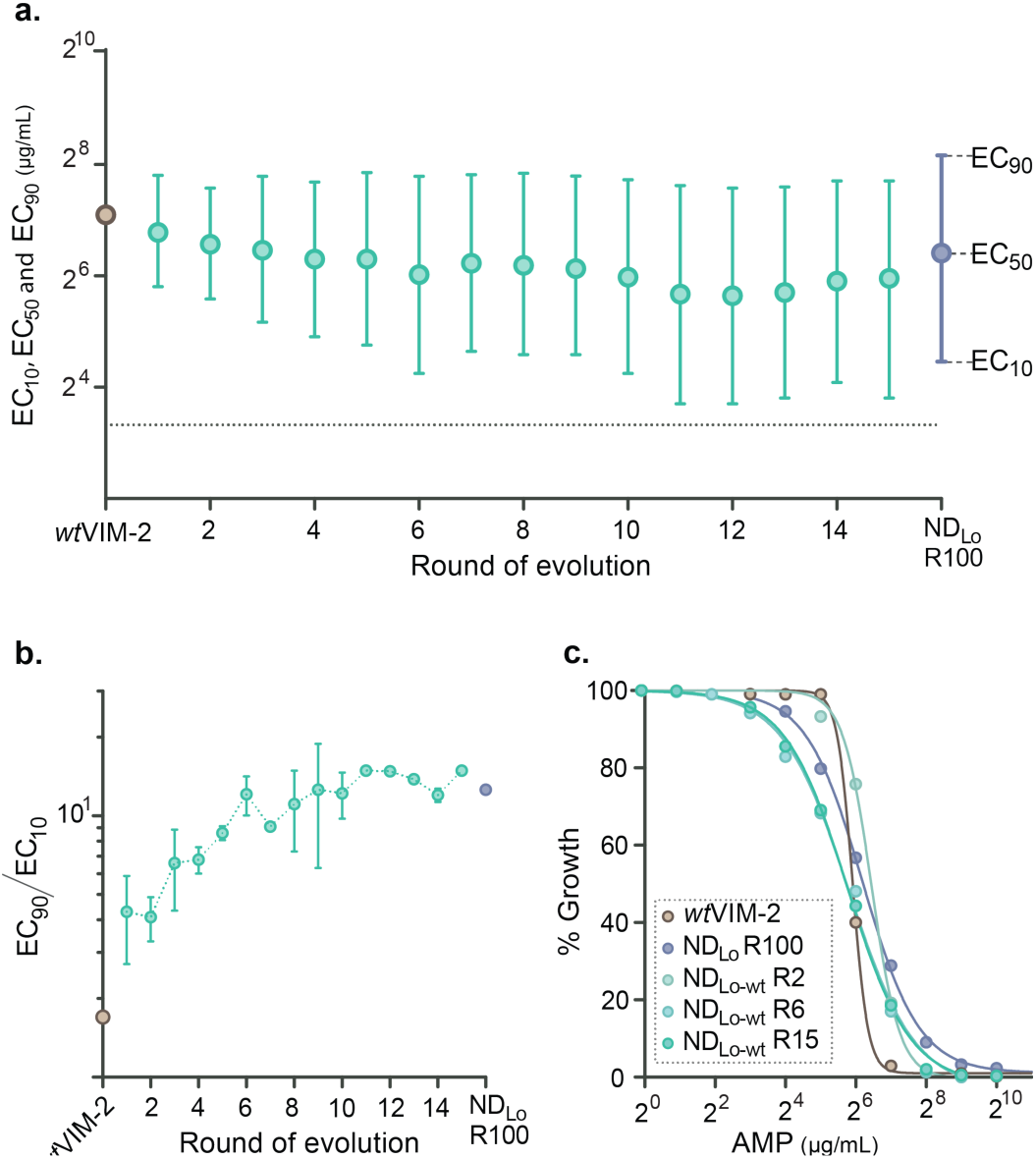
Phenotypic characterization of the NDLo-wt libraries. **a,** Changes in the ampicillin EC10, EC50 and EC90 of NDLo-wt over the experimental evolution. The central dot represents EC50, and bars at both ends for EC10 and EC90 respectively. **b,** Changes in EC90/EC10 during the evolution, error bars represent standard error between 2-4 biological replicates each NDLo-wt population. **c**, Ampicillin dose-response curves of select NDLo-wt libraries compared to NDLo and *wt*VIM-2. The data of **b** and **c** is shown in **Supplementary Table 4**.

### Neutral drift with threshold selection is the mechanistic basis for high phenotypic variation

Finally, we sought to determine how the intrinsic dynamics in neutral drift itself can explain observed high phenotypic variation. To this end, we developed a model which simulates the evolution of an antibiotic resistance gene, and evaluated how the evolution shapes phenotypic variation in a static environment (**Fig. 5**). Dynamics in neutral drift can be recapitulated as mutation-selection balance, in which the influx of different types of mutations (*e.g*., beneficial, neutral, deleterious mutations) is constantly shaped by selection to reach an equilibrium state^31–33^. In this equilibrium, phenotypic variation can arise when variants with different antibiotic resistance levels exhibit similar fitness levels and selectively neutral. A key attribute of our model compared to previous studies of mutation-selection balance is the threshold-like relationship between fitness and phenotype, where a selectively neutral zone exists above the selection threshold. Therefore, while previous models explored mutational effects directly on organismal fitness and resulting genetic diversity, we first consider mutations as effects on the protein phenotype at the molecular level, *i.e*., mutations may increase and decrease the resistance levels, and then evaluate the effect of phenotypic changes on the organismal fitness at a given antibiotic selection pressure. This allows for a more realistic model of protein evolution based on biochemical principles. We calculate the fitness of each variant using **equation (1)**:

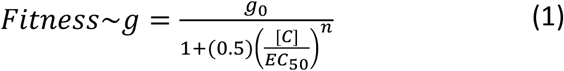

where *g*_0_ denotes the growth rate in the absence of ampicillin and *n* represents the Hill coefficient, [*C*] is the concentration of the antibiotic, EC_50_, the resistance level of the variant. The threshold-like fitness-phenotype relationship has been observed before in many proteins including *wt*VIM-2 (**Extended Data Fig. 7a**)^18,25,26^. In our model, the influx of mutations and variants within the population is constantly shaped by selection; variants can be selected and purged at rates proportional to their selection coefficients [**equation (2),** where *F* is fitness and *s* is the selection coefficient of the variant over *wt*VIM-2]:

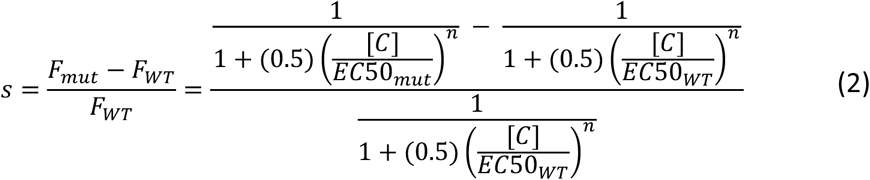

As mutations accumulate, variants with lower resistance levels than the antibiotic threshold will be purged, and variants retaining resistance above the threshold will survive. Importantly, a threshold relationship between fitness and resistance implies that there is a fitness plateau^26^, which comprises a selectively neutral zone for a range of resistance; variants with much higher resistance than the threshold would exhibit the same fitness level as variants with marginal resistance (**Extended Data Fig. 7a**).

**Fig. 5.**
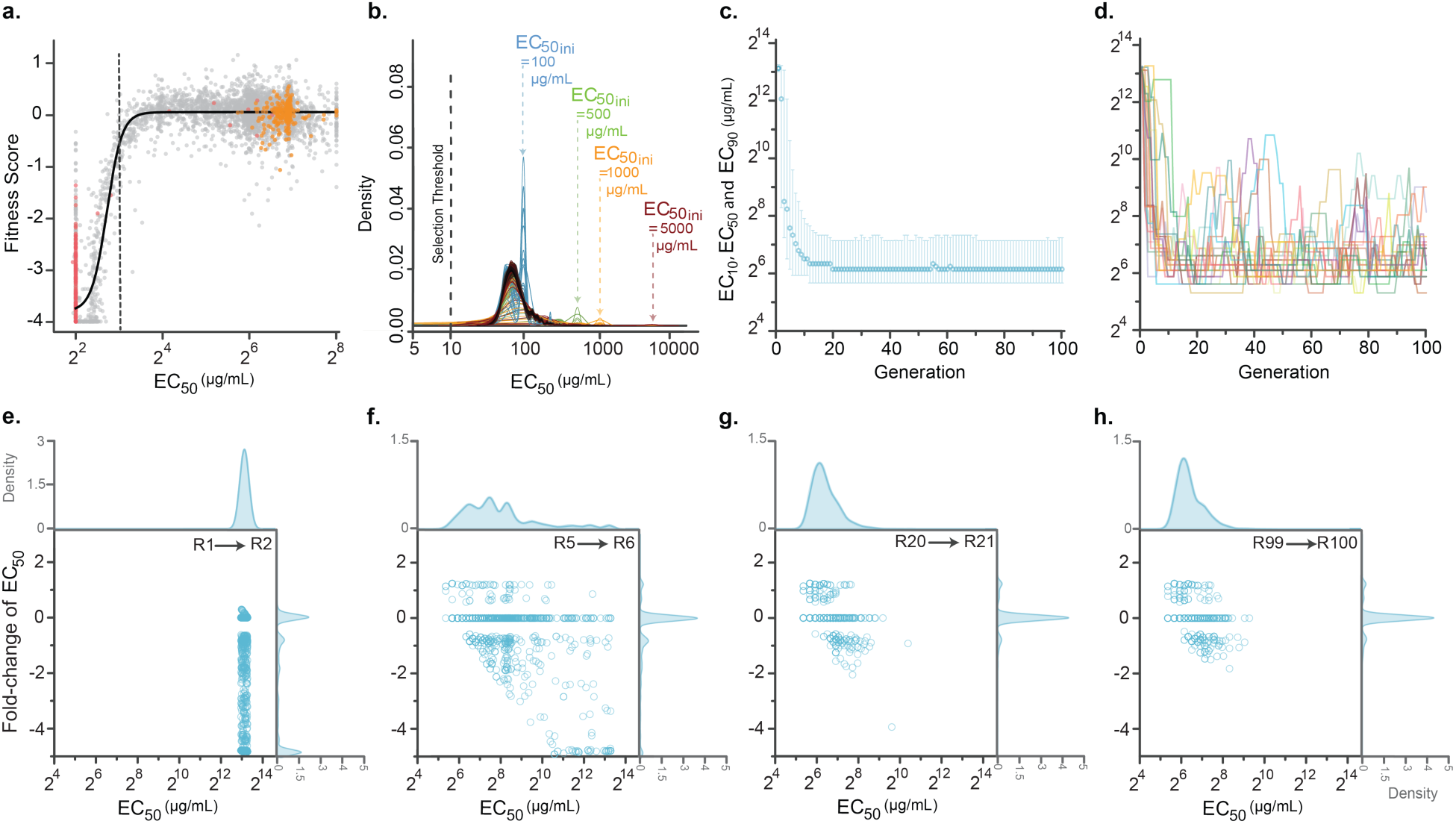
DME of *wt*VIM-2 and simulation of the NDLo trajectory. **a,** Resistance phenotype–fitness landscape of *wt*VIM-2 DMS library in the presence of AMP. Nonsense mutations are annotated in red, synonymous in orange and missense mutations in grey. The solid black curve indicates the line of best fit for a sigmoidal curve fit (equation (1)), while the black vertical dashed line indicates the AMP concentration used during selection (8µg/mL). Adapted *from Chen et al., 202*1^26^. **b,** Convergence of the resistance level of the NDLo populations starting from different initial conditions. The density of the ampicillin resistance level of the VIM-2 populations evolving under the NDLo regime is plotted as they evolve to the equilibrium state where different types of mutations are at balance. The trajectories for simulated evolution were started from variants with the initial EC50=100 µg/ml (blue, reflects the experimental NDLo-wt trajectory), 500 µg/ml (green), 1,000 µg/ml (orange), and 5,000 µg/ml (red, reflects the experimental NDLo trajectory). These initial populations converged to a final distribution of resistance levels shown in black. Simulation data can be found in **Supplementary Data 5**. **c,** Ampicillin EC10, EC50 and EC90 (µg/mL) of variants evolving under the NDLo regime across 100 rounds of simulated evolution, with an effective population size (Neff) of 10^4^. The selection threshold is indicated with the grey dotted line. **d,** Ampicillin EC50 values of 20 randomly picked variants from the simulated NDLo trajectory across 100 rounds (or generations) of evolution. **e.** Scatter plot of the distribution of mutational effects on the ampicillin EC50 of variants evolving under the simulated NDLo regime, compared to the ampicillin EC50 of the 10^3^ variants before acquiring the mutation. The mutational effects on resistance level are shown for the simulated variants from R1, R5, R20, and R99, as they move onto R2, R6, R21 and R100, respectively (the same data for additional rounds is given in **Extended Data Fig. 9**). Data and calculations for panels **c, d** can be found in **Supplementary Data 6**.

In order to conduct the most realistic simulations, we used the key parameters for the model using equation 1; *n* and [*C*] were empirically obtained from a global fitness-phenotype-environment landscape in our previous DMS study of VIM-2 (**Extended Data Fig. 7a** and **Methods**)^25,26^. We also utilized the experimentally obtained distribution of mutational effect (DME) on the level of antibiotic resistance (EC_50_) of *wt*VIM-2. The DME of *wt*VIM-2 represents largely 2% positive, 35% neutral and 63% negative mutations in terms of EC_50_ **(Extended Data Fig. 7b**).^25^ We performed the simulations with a population size of 10^4^, beginning from various EC_50_ starting points. At each round, the population experienced an influx of random mutations based on the DME with a rate of one mutation per variant per generation. A variant was propagated in the next cycle if it satisfied the probability of fixation using the equation described in **Methods**. We used a fixed distribution of DME in terms of fold change in EC_50_ during the simulations. The simulations strongly corroborated the characteristic features of the ND_Lo_ populations *i.e*., the populations reach identical and high phenotypic variation regardless of the resistance phenotype of the starting populations (**Fig. 5b**).

Next, we tracked mutational dynamics and their effects on fitness and EC_50_ during the simulations and identified several consequences for the equilibrium state in the distribution of phenotype (**Fig. 5b-h**, and **Extended Data Fig. 8**). When the starting population exhibits much higher resistance to the threshold, the population will accumulate more negative mutations, and consequently, the average resistance will decrease (**Fig. 5c-f**, **Extended Data Fig. 8a**, and **Extended Data Fig. 9**). However, as the average resistance approaches the threshold, fixation mainly occurs among positive, neutral, and mildly negative mutations while highly negative mutations are consistently purged out (**Fig 5c, f-h**, **Extended Data Fig. 8a**, and **Extended Data Fig. 9**). At mutation-selection balance, ∼10% of fixated mutations that changed the EC_50_ significantly, did so by increasing EC_50_ more than two-fold (**Fig. 5g-h**, and **Extended Data Fig. 9**) At this dynamic stage, high-resistance variants can sporadically emerge in the neutral zone without a substantial selective advantage and by stochastically acquiring positive mutations (**Fig. 5d-h,** and **Extended Data Fig. 9**). Such high-resistance variants can further accumulate mildly negative mutations which reduce their resistance levels. The sporadic emergence and disappearance of high-resistance variants create an equilibrium, resulting in observed phenotypic diversity.

We further found that the fraction of positive mutations in the DME establishes an upper bound to the resistance level of the population (**Extended Data Fig. 10**). Our simulations with low or no fraction of positive mutations in DME resulted in low phenotypic variation with steep declines in high-resistance variants, and mutation-selection balance work to largely eliminate negative mutations, while still accumulating neutral mutations. This is likely the case in the ND_Hi_ populations, as the population was drifting with an extremely high selection threshold for antibiotic resistance that exhausted the pool of positive mutations. Taken together, our results show that populations that evolve under mutation-selection balance and on a threshold-like fitness landscape can generate and maintain the phenotypic variation. The level of such variation for a given population size is dictated by the DME and the shape of the phenotype-fitness relationship (**Extended Data Fig. 7**).

## Discussion

Using experimental evolution, we demonstrate that variation in phenotype (antibiotic resistance level) of VIM-2 β-lactamase can be simply promoted through neutral drift, *i.e*., evolution in a static environment. Our observations provide important implications for understanding and predicting the evolution of antibiotic and drug-resistance genes in the environment and clinics^30,34^. Our results suggest that the emergence of high antibiotic resistance pathogens can be promoted even in the presence of trace amounts of antibiotics, such as concentrations observed in the environment due to global anthropogenic antibiotic contaminations^35–37^. Such “hidden” high-resistance variants within the population can serve as a springboard to readily adapt when the concentration of antibiotics is increased, *e.g*., applying antibiotics to patients, animals, and environments. Moreover, as the ND_Lo_ populations exhibited variable and higher substrate specificity against non-selected β-lactams, the population can encompass further evolvability against other antibiotics as well. Further understanding such evolutionary dynamics in natural environments would be critical for combatting the emergence and dissemination of multidrug-resistant pathogens.

More generally, our findings provide a new and robust mechanistic explanation for the universal existence of phenotypic variation and adaptive capacity within evolving populations. Our observations explain how phenotypic variation of a trait which is directly under selection pressure can be generated and maintained through evolution in a static environment. Indeed, many natural proteins evolve via neutral drift, *i.e.*, under purifying selection pressure to maintain their function above a certain threshold^38^. Also, the threshold-like relationship between the traits of protein (function and stability) and fitness is commonly described and even considered as a universal attribute^17,39–42^. Furthermore, many experiments showed the existence of a considerable fraction of positive mutations to enhance protein function and stability^18,25,26,43,44^, indicating that the selection thresholds for many proteins in nature may be modest and more similar to the ND_Lo_ compared to ND_Hi_^45,46^. In consequence, many, if not most, biological molecules (and by extent, organisms) will inevitably exhibit phenotypic variation within a population and species. Importantly, the mechanism we found in this study are not incompatible with previously described mechanisms for phenotypic variation such as differential selection and environmental perturbations, as these mechanisms simply add more phenotypic variation in the population. Moreover, variants with much higher functional levels than the selection threshold can emerge and be maintained during neutral drift in a simple and static environment without being selected for. Thus, neutral drift under threshold selection plays a key role in facilitating the evolutionary capacity to adapt to environmental perturbations.

## Acknowledgments

We thank members of the Tokuriki lab for the discussions and comments.

## Funding

We thank the Canadian Institute of Health Research (CIHR) Project Grants (AWD-019305 and AWD-018386) for the financial support.

## Author contributions

Conceptualization: NT, ANE, PD, RDS, AWRS

Methodology: NT, ANE, PD, RDS, LK, RJ, BEL

Investigation: NT, ANE, PD, RDS, JZC, AWRS

Visualization: ANE, PD, JZC

Funding acquisition: NT

Project administration: ANE, RDS, LK, RJ, BEL

Supervision: NT, RDS, PD, AWRS, JZC

Writing – original draft: ANE, NT, PD, RDS

Writing – review & editing: ANE, NT, PD, RDS, JZC, AWRS

## Conflict of Interest

Authors declare no competing interests.

## Data Availability

All experimental data can be provided upon request from the corresponding author.

Description of supplementary data file contents are given below.

***Supplementary Data 1:*** Adaptive walk ampicillin selection regime and number of surviving variants after selection.

***Supplementary Data 2:*** Genotypic analysis of individual variants from the AW, ND_Lo,_ ND_Hi_ and ND_Lo-wt_ libraries and their β-lactam antibiotic MIC.*

***Supplementary Data 3:*** β-lactam MIC values of variants from the AW, ND_Lo,_ ND_Hi_ libraries and analyses.

***Supplementary Data 4:*** Ampicillin dose-response curve data of AW, ND_Lo,_ ND_Hi_, ND_Lo-wt_, P0-ND_Lo,_-R100, P10-ND_Lo,_-R100, and P10-ND_Lo,_-R60 libraries.

***Supplementary Data 5:*** Raw ampicillin EC_50_ values of 100 variants from simulated ND_Lo_ evolutionary trajectory with different starting ampicillin EC_50_ values across 100 generations.

***Supplementary Data 6:*** Raw ampicillin EC_50_ and fitness values of 100 variants from simulated ND_Lo_ evolutionary trajectory across 100 generations.

***Supplementary Data 7:*** Relative fraction of mutations that affect enzyme phenotype (*ie.* ampicillin EC_50_; positive, negative and neutral) and the fitness of variants from simulated ND_Lo_ evolutionary trajectory across 100 generations per generation.

***Supplementary Data 8:*** Fasta files of the DNA sequences of the cloning plasmid pIDR with *wt*VIM-2 gene inserted into the cloning region of the randomly-picked mutant VIM-2 variants.

*The MIC values are not present for all β-lactams used here for screening (meropenem, cefotaxime, ampicillin) for all mutant VIM-2 variants.

## Statistical Analysis and Code Availability

We used R (v4.2) within Jupyter Notebook (v6.5.2) for all statistical analyses of the simulations. In particular, we plotted the density of EC_50_ values in Figure 5b using the kernel density function in R. The code to simulate the evolution of ND_Lo_ is available at the GitHub repository: https://github.com/dasmeh/Neutral_Zone/.

## Materials and Methods

### Construction of VIM-2 libraries for the AW and ND_Hi/Lo/Lo-wt_ trajectories

The wild-type (wt) VIM-2 gene including its signal peptide sequence was synthesized (Bio Basic Inc.) and subcloned into a low-copy number plasmid, pIDR2, under a constitutive TEM-1 derived promoter and a chloramphenicol resistance marker (*cat*), using NcoI and XhoI restriction enzyme sites. For the AW trajectory, randomly mutagenized libraries of wt VIM-2 were created by error-prone PCR with the nucleotide analogues, 8-oxo-2’-deoxyguanosine-5’-triphosphate (8-oxo-dGTP) and 2’-deoxy-P-nucleoside-5’-triphosphate (dPTP) (TriLink). For each library, two independent PCRs with 8-oxo-dGP and dPTP were performed to ensure a balance between transition versus transversion type nucleotide mutations and a specific mutation rate. Each 25 μL PCR consisted of 1 x GoTaq Buffer (Promega), 3 μM MgCl2, 0.1 μM of each primer, 0.2 mM of dNTPs, 1.00 U of GoTaq DNA polymerase (Promega), 1 ng of template plasmid, and either 100 μM of 8-oxo-dGTP or 1 μM of dPTP. The first PCR was programmed as follows: an initial denaturation (95°C for 2 minutes), followed by 20 cycles of 95°C for 30 seconds, 58°C for 60 seconds, 72°C for 60 seconds, before a final extension step (72°C for 3 minutes). The PCR products were then subsequently purified with the EZ.N.A.® Cycle Pure PCR Purification Kit (OMEGA Bio-tek Inc), quantified, mixed and used as a template for the second PCR. Each 50 μL amplification PCR consisted of 1 x GoTaq Buffer (Promega), 3 μM MgCl2, 0.1 μM of each primer, 0.25 mM of dNTPs, 1.0 U of GoTaq DNA polymerase (Promega), and 5 ng of each PCR product from the two previous reactions. The reaction was run as the first PCR but with 35 cycles. The PCR products were purified with the EZNA Cycle Pure PCR Purification Kit, digested with NcoI (FastDigest, ThermoFisher Scientific™) and XhoI (FastDigest, ThermoFisher Scientific™) for the AW and ND_Lo_ and ND_Hi_ libraries and NcoI and KpnI (FastDigest, ThermoFisher Scientific™) for ND_Lo_-wt libraries for a 1 hr at 37°C. The pIDR plasmid was also digested by NcoI and XhoI, or NcoI and KpnI for 3 hours at 37°C. The digested plasmid was subsequently purified from 1% agarose gel using gel purification columns, the digested PCR products were purified with the E.Z.N.A.® Cycle Pure PCR Purification Kit. The ligation mixture (10 μL) consisted of 1 × T4 DNA ligase buffer (ThermoFisher Scientific™), 5 U of T4 DNA ligase (ThermoFisher Scientific™), 10-8 ng of prepared vector, and 30-40 ng of prepared mutagenized insert, was incubated at room temperature for 1 hour. The ligations were then purified with a MicroElute kit (OMEGA Bio-tek Inc.) and eluted with 20 μL of water.

### Selection of the libraries in the presence ampicillin

To create the AW and ND_Hi/Lo/Lo-wt_ libraries, 4-5 μL of the purified ligation mixtures were transformed to *E.cloni*® 10G *E. coli* cells (Lucigen Corp.) using electroporation. For the AW trajectory, the transformants were grown overnight at 30°C in 10 mL of LB media supplemented with 34 μg/mL of chloramphenicol. Then, 100 μL of a 1:100 dilution of the overnight culture was plated onto a series of LB agar plates containing 2-fold increases in the concentration of ampicillin from 2 to 8192 μg/mL. The plate with the highest concentration of ampicillin with colony counts between 100 and 1,000 colonies was collected (**Supplementary Data 1**). The plasmids were extracted from the colonies and used as the template for the next round. For the ND_Hi/Lo/Lo-wt_ trajectories, the transformants were plated onto large LB agar plates containing 34 μg/mL of chloramphenicol in addition to 1,000 μg/mL of ampicillin for the ND_Hi_ libraries, and 10 μg/mL of ampicillin for the ND_Lo/Lo-wt_ libraries.

### Antibiotic dose-response curves to calculate EC_50_ and EC_90_/EC_10_

To assess the average resistance level (EC_50_; or effective concentration that inhibits the growth of 50% of the population) and the diversity of the resistance levels (approximated by the ratio of EC_90_/EC_10_) present within each population, antibiotic dose-response curves were obtained using cell culture assays using 96-well plates. First, glycerol stocks of *E.cloni*® 10G *E. coli* cells harboring single libraries were inoculated in 3 mL LB media supplemented with 34 μg/mL chloramphenicol (LB-Cm) for 16 hours at 30°C. The OD_600_ value of these cultures was calculated, and diluted to have an OD_600_ value of 0.0015 in a 10 mL day culture. The cultures were then grown for 1 hour 20 minutes at 30°C, or until the OD_600_ of cultures reached 0.01-0.02. 180 μL of the day cultures were mixed with 20 μL LB-Cm either containing no additional antibiotics or supplemented with 11 different concentrations of antibiotic with 2-fold increments of ampicillin/cefotaxime/meropenem, in 96-well assay plates (ThermoFisherScientific™ CORNING). The cultures were grown for 6 hours at 37°C, and OD_600_ of the cultures were measured. The ‘percent survivals’ of each library at each β-lactam antibiotic concentration was calculated comparing OD_600_ value of *E. coli* cultures in the absence and presence of antibiotics in the culture. The values were then fit onto a Hill equation (1) with a top constraint of 100 using the PRISM™ 9.0 software.

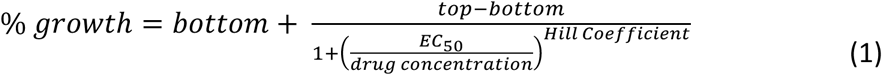

The EC_50_, EC_90_ and EC_10_ values of the curve were extracted to calculate EC_50_ and EC_90_/EC_10_. Each assay was carried out with two technical replicates (two independent culture in the same 96 well plate), and at least 2 biological replicates (the same experiment on different days) were carried out for each library.

### Measuring the minimum inhibitory concentration of individual variants

To quantify the ampicillin resistance level conferred by individual variants from VIM-2 libraries to *E. coli*, we used agar-plate based assays to determine the minimum inhibitory concentration (MIC) of antibiotics to *E. coli* carrying a VIM-2 variant. *E. coli* cells harboring a single VIM-2 variant were grown in 500 μL LB-Cm at 30°C overnight in a deep-96-well plate. The next day, 5 μL of the overnight culture was inoculated into 195 μL of LB-Cm in quadruplicate in a standard 96-well plate and grown for 3 hours at 37°C. The cultures were then plated with 96-well replicator pins on a series of 15 mm LB agar plates with increasing levels of antibiotics (two-fold increases in ampicillin, meropenem, and cefotaxime from, 2 to 32,768 μg/mL, 0.016 to 64 μg/mL, and mL 0.032 to 4096 μg/mL respectively). The agar plates were subsequently incubated overnight at 37°C. The next day, the MIC was determined by identifying the concentration of antibiotics by which no growth was observed in at least three of the four replicates for each variant. Descriptive Statistics on the data was conducted on PRISM 9.0. The specificity coefficients were calculated using the following formula:

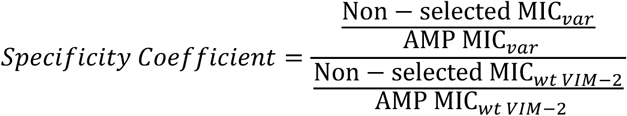

### Sequencing of individual variants

24-96 single colonies were randomly picked from selected libraries, and the VIM-2 gene region of the plasmid was PCR amplified using NEB Taq2x Master Mix using the manufacturer’s protocol, with an initial denaturation (95°C for 2 minutes), followed by 30 cycles of denaturation (95°C for 30 seconds, 58°C for 60 seconds and 72°C for 60 seconds), before a final extension step (72°C for 3 minutes). The PCR products were then purified enzymatically by treatment with ExoI (ThermoFisherScientific™) and FastAP (ThermoFisherScientific™) for 1 hour at 37°C, and then the enzymes were inactivated via heat treatment of the sample by incubation at 85°C for 15 minutes. The purified products were sent for Sanger sequencing (Azenta™). The sequence results were visually inspected in Geneious® bioinformatics software and the mutations were identified by comparing each mutant VIM-2 gene sequence to the *wt*VIM-2 gene sequence. The identified amino acid and nucleotide mutations were used to calculate the percent identity each variant shared with *wt*VIM-2 to determine divergence from the *wt*VIM-2 sequence, and the percent identity shared by each variant with other variants from the same library, to calculate within library diversity. Any insertion or deletion variants were brought to the same length as *wt*VIM-2 by adding ‘X’ in place of the deleted residues (deletion only occurred at the last 5-10 amino acid residues), and removing any insertion mutations at the 3’ end of the gene due to the mutations randomly introduced to the stop codon.

### Calculation of random walk threshold for N_a_/N_t_

To estimate the expected N_a_/N_t_ ratio from a completely random accumulation of mutations (i.e., all mutations are tolerated), we first calculated the N_a_/N_t_ ratio for each codon when mutated to all other codons. N_a_ = 1 if the codons encode different amino acids, else N_a_ = 0. Nt is simply the number of nucleotide differences between codons, regardless of the exact base. We then take the average N_a_/N_t_ of all possible mutations for each codon to get the random walk N_a_/N_t_ for a given codon. To estimate the N_a_/N_t_ for a drifting sequence starting from wtVIM-2, we calculate an average of all random walk N_a_/N_t_ ratios for the codons, weighted by the frequency of each codon in the *wt*VIM-2 sequence, arriving at a final value of 0.46.

### Phylogenetic tree construction

656 individual sequences of randomly picked variants from the AW, ND_Lo,_ and ND_Hi_ libraries were subjected to multiple sequence alignment (MSA) using ClustalW. The MSA was used to construct a phylogenetic tree using IQ-TREE Multicore Version 2.12 COVID-edition (March 30^th^ 2021) and with an ‘ultrafast_bootstrap’ bootstrap replicate number of 5,000. The tree was visualized and annotated using the ITOL™ tool.

### Population genetic simulations to capture evolutionary dynamics in the trajectories

The distribution of mutational effects (DMEs) of *wt*VIM-2 for ampicillin resistance (EC_50_) was experimentally obtained in our previous deep mutational scanning (DMS) study of VIM-2 (**Supp. Fig. 1**)^25,26^We also obtained the relationship between EC_50_ and the fitness of *E. coli* harbouring VIM-2 variants.

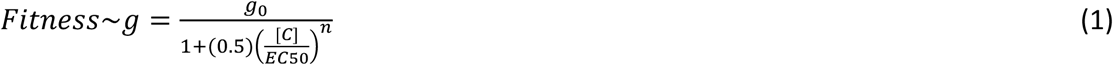

where, average fitness of mutant VIM-2 variants was estimated by the calculation of the bacterial growth rate (*g*) was modelled by a Hill function of the MIC concentration of ampicillin, the relative growth rate in the absence of ampicillin (*g_0_*) the Hill-coefficient (*n*), the concentration of the antibiotic at which selection was performed [C], and finally the concentration at which 50% of growth of a single population was inhibited (EC_50_). We used the parameters of Equation 1 from our previous study in which we determined the empirical fitness landscape of VIM-2 populations (**Supp. Fig. 2a**)^26^. In this previous study, using the correlations between the fitness scores of each single point mutant of *wt*VIM-2 under ampicillin selection with the EC_50_ level of each single point mutant. The dose-sensitivity parameter Hill-coefficient (*n*) was estimated to be 5. Using the DME of *wt*VIM-2 and the empirically obtained the fitness-phenotype-environment relationships, population genetic simulations reflecting neutral drift experiments VIM-2 variants were conducted to obtain the physiological fitness landscape of antibiotic resistance in our model system.

We first divided the phenotype space of EC_50_ values into *n=* 1,000 different states, ranging from EC_50_=0 to EC_50_=10,000. We assumed that a VIM-2 variant occupies one of these states and can transition from one to another by single point mutations. We then took an approach akin to Discrete Time Markov Chain simulations to simulate the evolutionary trajectory of VIM-2 in our experiments^47^. We constructed a 1,000 by 1,000 matrix that represents the transition probability matrix for transitions between different states and simulated the evolution of 1,000 Markov chains, each representing a VIM-2 variant in the population. All these Markov chains were initially in the same state and evolved to other states according to our transition probabilities.

The transition probability between each two states (i, and j) is the product of two probabilities, one that represents mutations, *P_mutation_*(*i*→*j*)and the other represents selection, *P_selection_*(*i*→*j*):

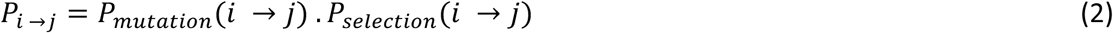

We calculated *P_mutation_*(*i*→*j*), i.e., the probability that VIM-2 transitions from the state *i*^th^ to *j*^th^ with single point mutations, from experimentally determined DME distribution. The second term, *P_selection_*(*i*→*j*), shows the fixation probability of such mutations. To estimate this probability, we related EC_50_ to fitness (Equation 1), and calculated the selection coefficient of any arising mutation on the wtVIM-2 background as:

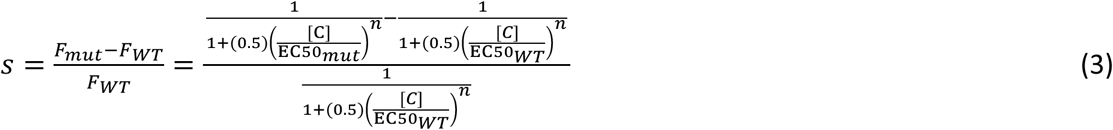

Here, [C] is the concentration of ampicillin in the media which takes the values of 10 µg/ml and 1,000 µg/ml, corresponding to the weak and the strong selection strengths used for the two experimental neutral drift trajectories, respectively. We then used the probability of fixation of arising mutations in a monoclonal haploid population from the Kimura formula^48^:

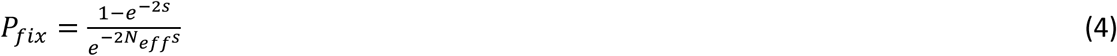

where N_eff_ is the effective population size and is 10^4^ in our simulations which is ∼ equal to the number of colonies sampled from LB-agar plates in each generation. We started the simulations with 10^4^ Markov chains all starting from a single state (*i.e*., initial EC_50_ value), and let these states evolve according to the transition probabilities and recorded their states after 100 rounds of mutation and selection (**Supp. Fig. 2b**).

**Supplementary Fig. 1.**
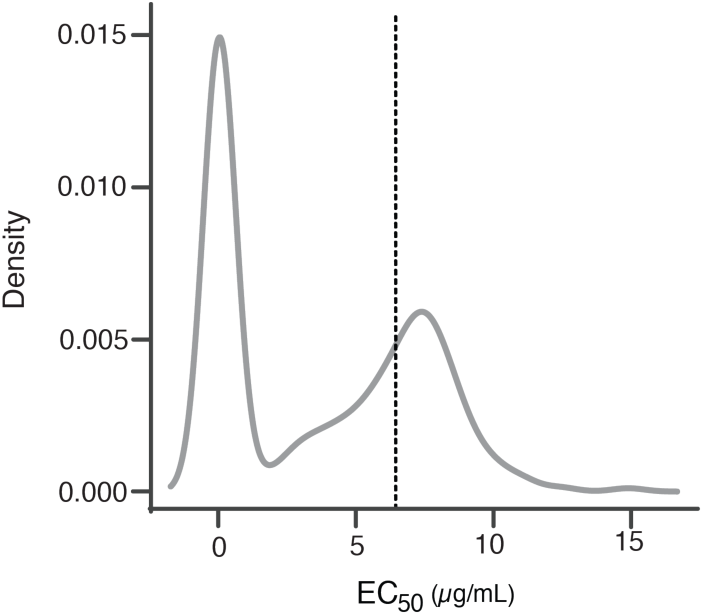
Distribution of the fitness effects of mutation on EC50 converted from DMS scores using Equation 1. The dashed line represents EC50 of the *wt*VIM-2.

We observed that simulation of ND_Lo_ trajectory shows an excellent agreement with the results obtained experimentally (**Supp. Fig. 2c**).

Moving forward, we also investigated the effect of increasing the selection strength (ampicillin concentration in experiments) on the evolutionary dynamics of our simulated VIM2 populations to understand the possible differences between ND_Lo_ and ND_Hi_ populations. To this aim, we systematically varied the ampicillin concentration in our simulations and generated evolutionary trajectories for populations with [AMP]=10, 100, 200, 300, 400, 500, 600, 700, 800, 900, and 1,000 μg/mL (**Supp. Fig. 2d**). For all these simulated populations, the mutation-selection balance was established within 15-20 mutations, in agreement with our observations in ND_Lo_ and ND_Hi_ populations.

**Supplementary Fig. 2.**
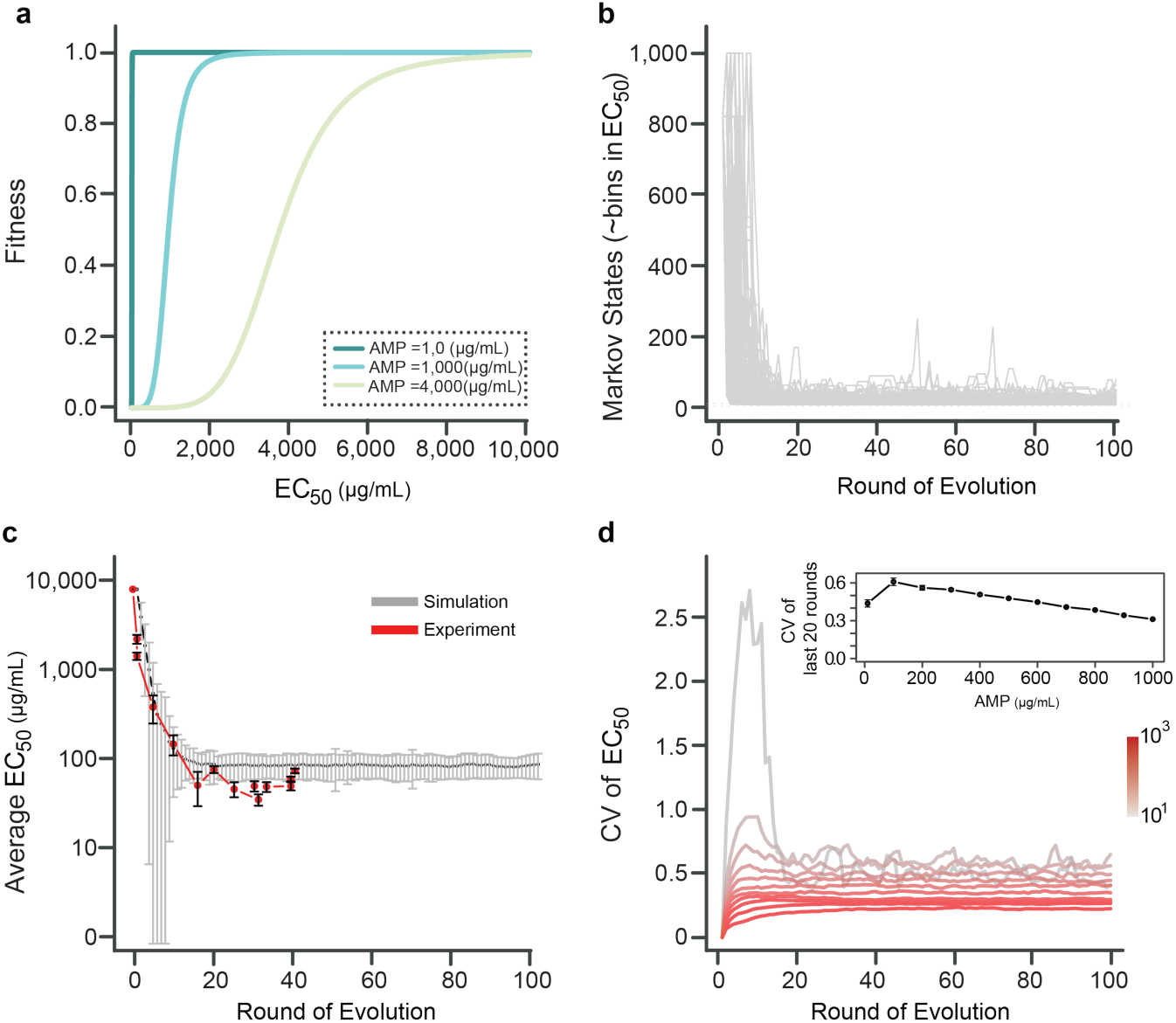
Simulation of the neutral drift versus experimental data. **a**, Fitness function relating growth rate to EC50 at ampicillin concentrations of 10, 1,000 and, 4,000 µg/ml shown in dark cyan, cyan and light green, respectively. **b**, The evolutionary dynamics of the NDLo population. Each individual VIM-2 protein transitions between different Markov states that corresponds to different EC50 bins by mutations. The probability of fixation is calculated from Equation 2. In this plot, all variants have an initial EC50 of 8192 µg/ml. **c,** Simulated (in gray) versus experimental (in red) average EC50 of the population as a function of the number of amino acid mutations. **d,** The coefficient of variation (CV) of the simulated population’s EC50 as a function of ampicillin concentration, [AMP]. We varied [AMP] from 10 to 1,000 µg/ml in simulations. The inset of panel d shows the average coefficient of variation of EC50 in the last 20 rounds of evolution for populations evolving under different ampicillin concentrations.

One interesting distinction between simulated populations at lower and higher ampicillin concentrations was the degree to which these trajectories were subject to neutral drift. As a result of the fitness function around the average MIC of the adapted population (∼11,000 µg/ml) neutral drift causes a decrease in EC_50_ proportional to the rate of fixation of neutral mutations (∼ 1/N_eff_) for populations that are subjected to selection at lower ampicillin concentrations. In contrast, the resistance fitness landscapes at higher ampicillin concentrations are more curved compared to lower concentrations, leading to a stronger selection pressure in the evolution of such populations (**Supp. Fig. 2a**). Therefore, upon the first mutational cycle both forces of mutation and selection more likely influence the MIC values of different variants in trajectories undergoing selection with higher ampicillin concentrations, compared to the trajectories with lower ampicillin concentrations which expectedly results in a lower coefficient of variation of phenotypic values. Indeed, the coefficient of variation of EC_50_ values decreased as the ampicillin concentration increased from 10 µg/ml to 1,000 µg/ml (**Supp. Fig. 2d**). This result is in full agreement with the observed differences between ND_Lo_ and ND_Hi_ (**Supp. Table 1, 2**). Altogether, neutral drift permits exploration of a wider range of fitness landscape at lower antibiotic selection thresholds.

## Extended Data Figures

**Extended Data Fig. 1.**
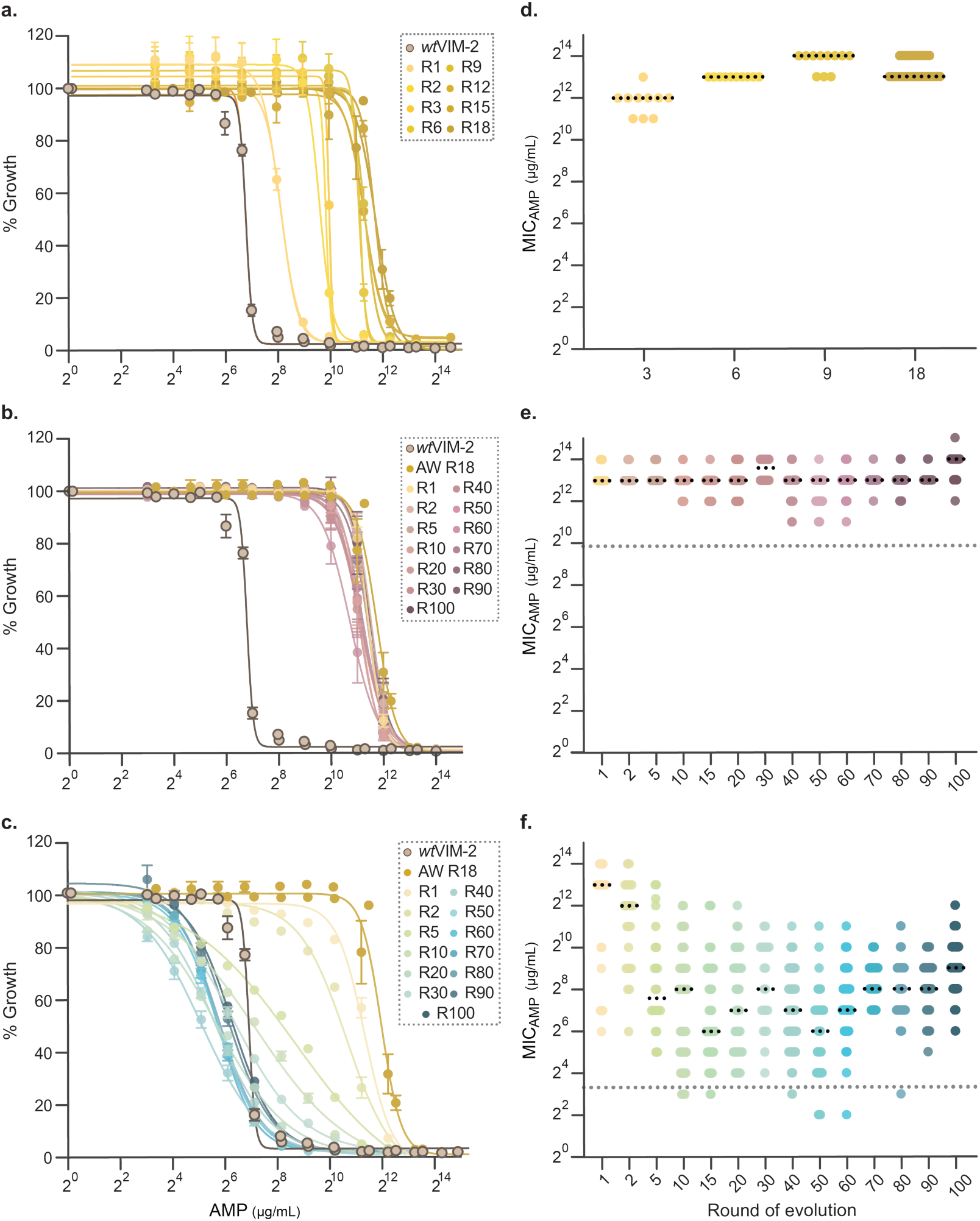
Population and individual level phenotypic analysis of the AW, NDHi and NDLo libraries. **a-c,** Ampicillin dose-response assay curves of select libraries from the AW (**a**), NDHi (**b**) and NDLo (**c**) libraries. Error bars depict standard error. At least 2 biological replicates were carried out for each DR assay. **d-f,** Distribution of ampicillin minimum inhibitory concentration (MICAMP) values of 24-96 individual variants randomly picked from the AW (**d**), NDHi (**e**), and NDLo (**f**) libraries. The average MICAMP of the libraries is shown with black dots, while the antibiotic selection threshold the libraries were subjected to is shown with grey dots. The data for the figures can be found in **Supplementary Data 3, 4.**

**Extended Data Fig. 2.**
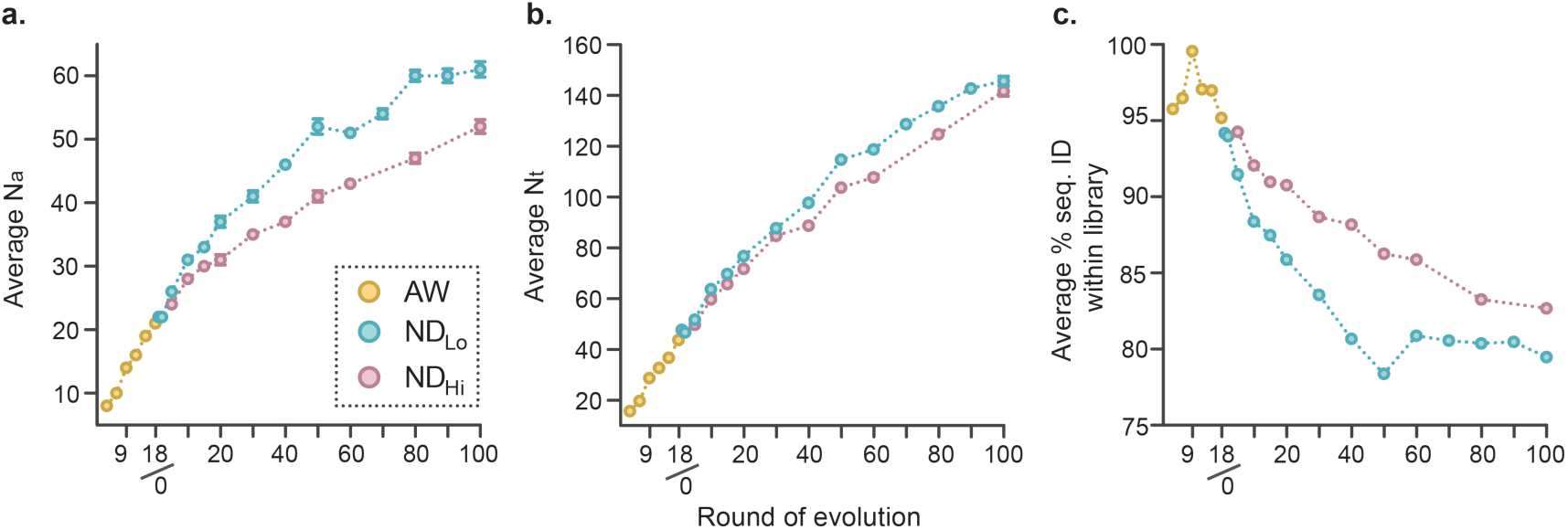
Genotypic analysis of the AW, NDHi and NDLo libraries. **a**, **b**, Average number of nucleotide mutations (Nt) (**a**) and amino acid mutations (Na) (**b**) present in the experimental evolution libraries, compared to *wt*VIM-2. Error bars depict standard error. **c**, Average percent sequence identity shared between the enzyme variants within the evolution libraries. The data for the figures are shown in **Supplementary Table 3**, and can be found in **Supplementary Data 2**.

**Extended Data Fig. 3.**
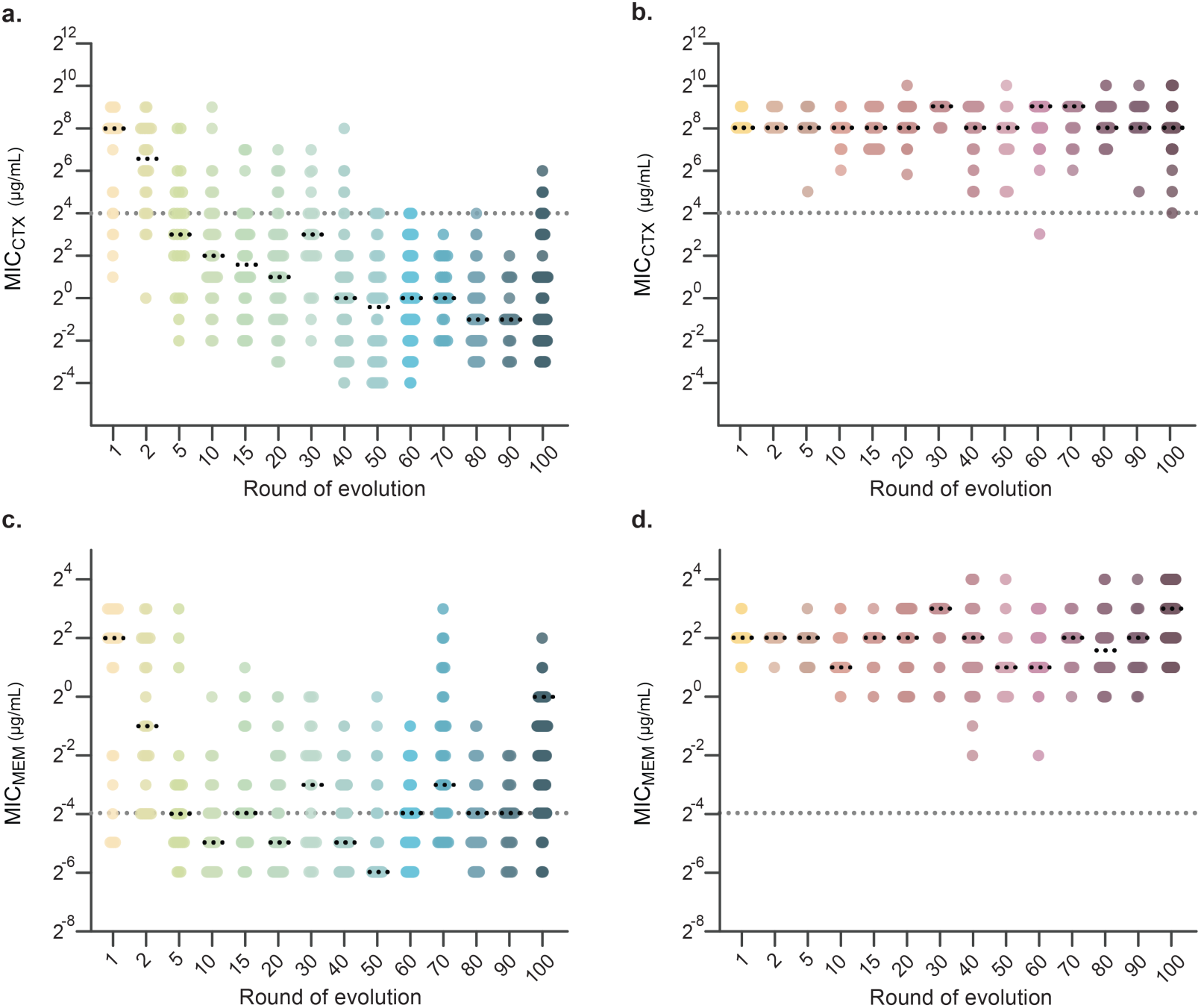
Phenotypic analysis of individual variants from the AW, NDHi and NDLo libraries for non-selected β-lactam antibiotics. Distribution of cefotaxime and meropenem minimum inhibitory concentration MIC values of 24-96 individual variants randomly picked from NDL (**a, c**) and NDHi (**b, d**) libraries, respectively. Mean MIC for each library is shown as black dots, and the *wt*VIM-2 resistance level for each antibiotic is shown as grey dots. The data for the figures can be found in **Supplementary Data 3.**

**Extended Data Fig. 4.**
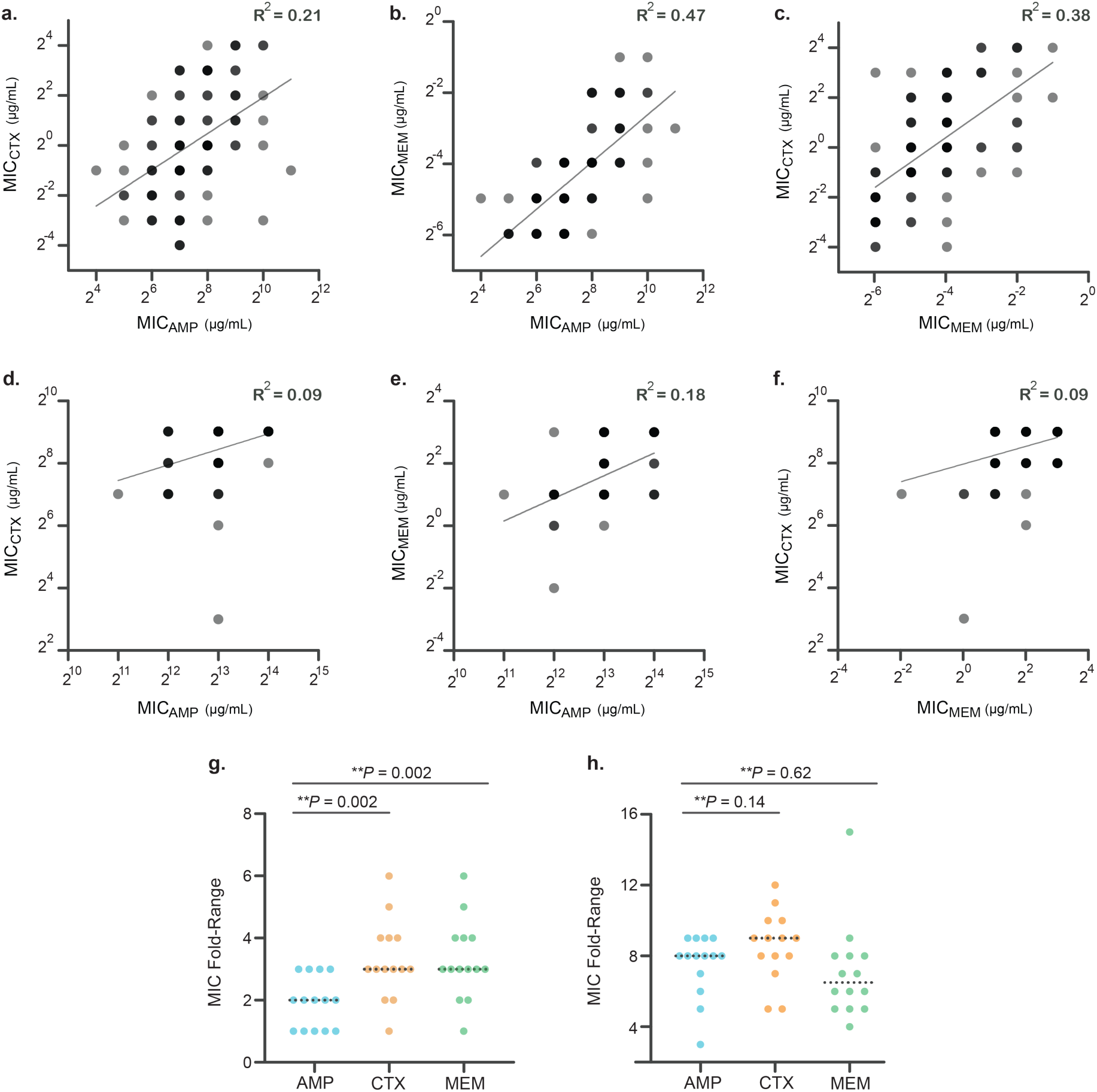
Comparisons of the MICs of individual variants for non-selected and selected β-lactam antibiotics. **a-f,** Correlation of the log2 of cefotaxime and ampicillin, meropenem and ampicillin, and cefotaxime and meropenem MICs of 91 individual variants from the NDLo (**a-c**) NDHi R60 (**d-f**) libraries, each (total *n*=182). The correlation coefficient for each pair is given on the top right of the graphs. Grey dots reflect resistance phenotypes observed in a single individual, while black dots reflect ones with >2 variants. **g-h.** Comparison of the fold-difference range of ampicillin (blue), cefotaxime (orange) and meropenem (green) MICs of 91 individual variants from the NDLo (**g**) NDHi (**h**) R60 libraries, each. *P*-values calculated via unpaired two-sided t-test on the 2-fold difference range of ampicillin and non-selected antibiotic MICs are shown on the top of the graphs. The data used for the figures can be found **in Supplementary Data 3.**

**Extended Data Fig. 5.**
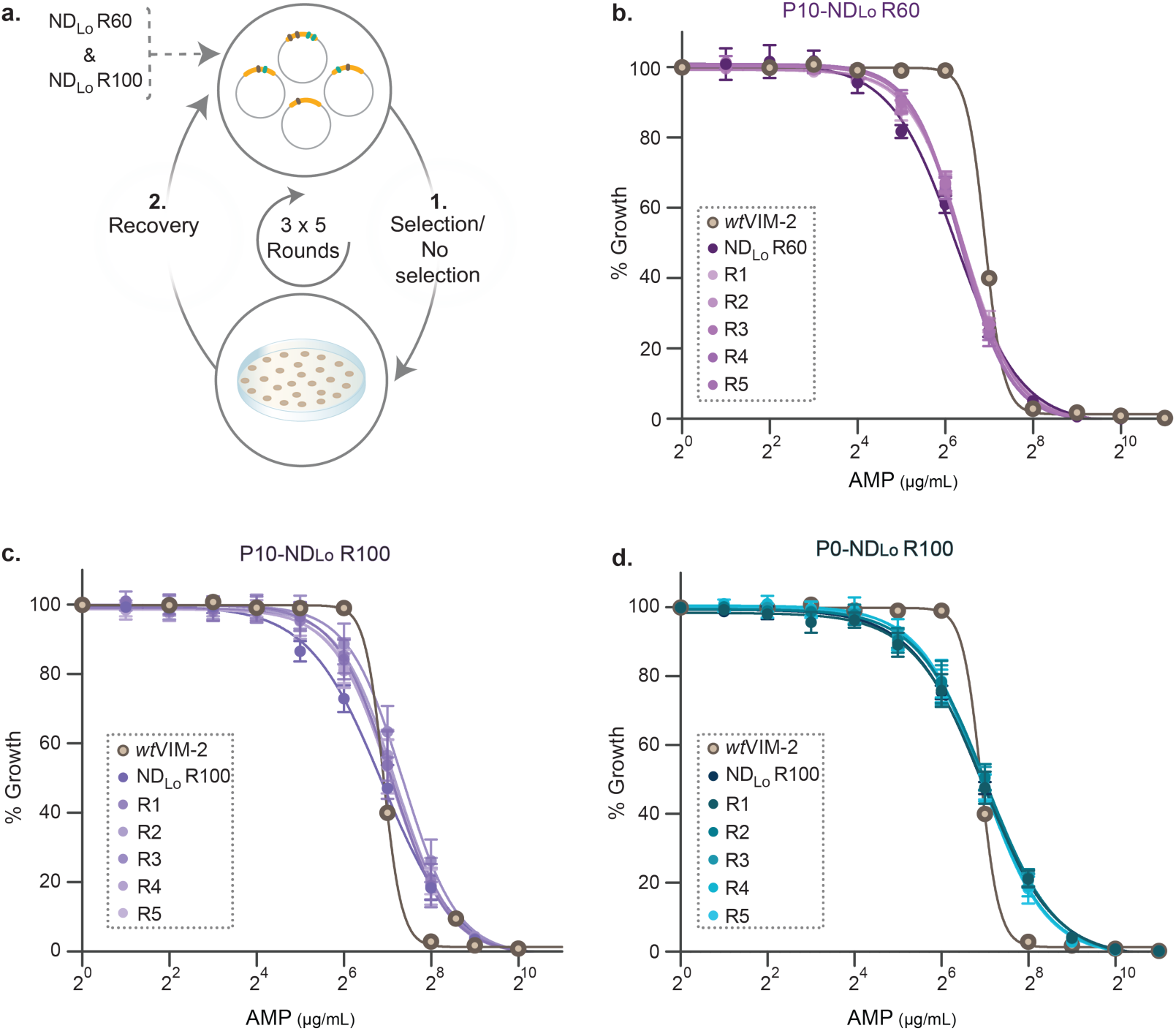
Scheme of passaging experiments and population-level phenotypic characteristics. **a,** Scheme depicting the passaging experiments of NDLo R60 and R100 libraries under selection by 10 µg/mL ampicillin (P10-NDLo R60 and P10-NDLo R100, respectively), and of NDLo R100 under no selection by ampicillin (P0-NDLo R100). Ampicillin dose-response assay curves of the five P10-NDLo R60 (**b**), P10-NDLo R100 (**c**), and P0-NDLo R100 (**d**) libraries. Error bars depict the standard deviation, two biological replicates were carried out for each library. The data for the figures are presented in **Supplementary Data 4.**

**Extended Data Fig. 6.**
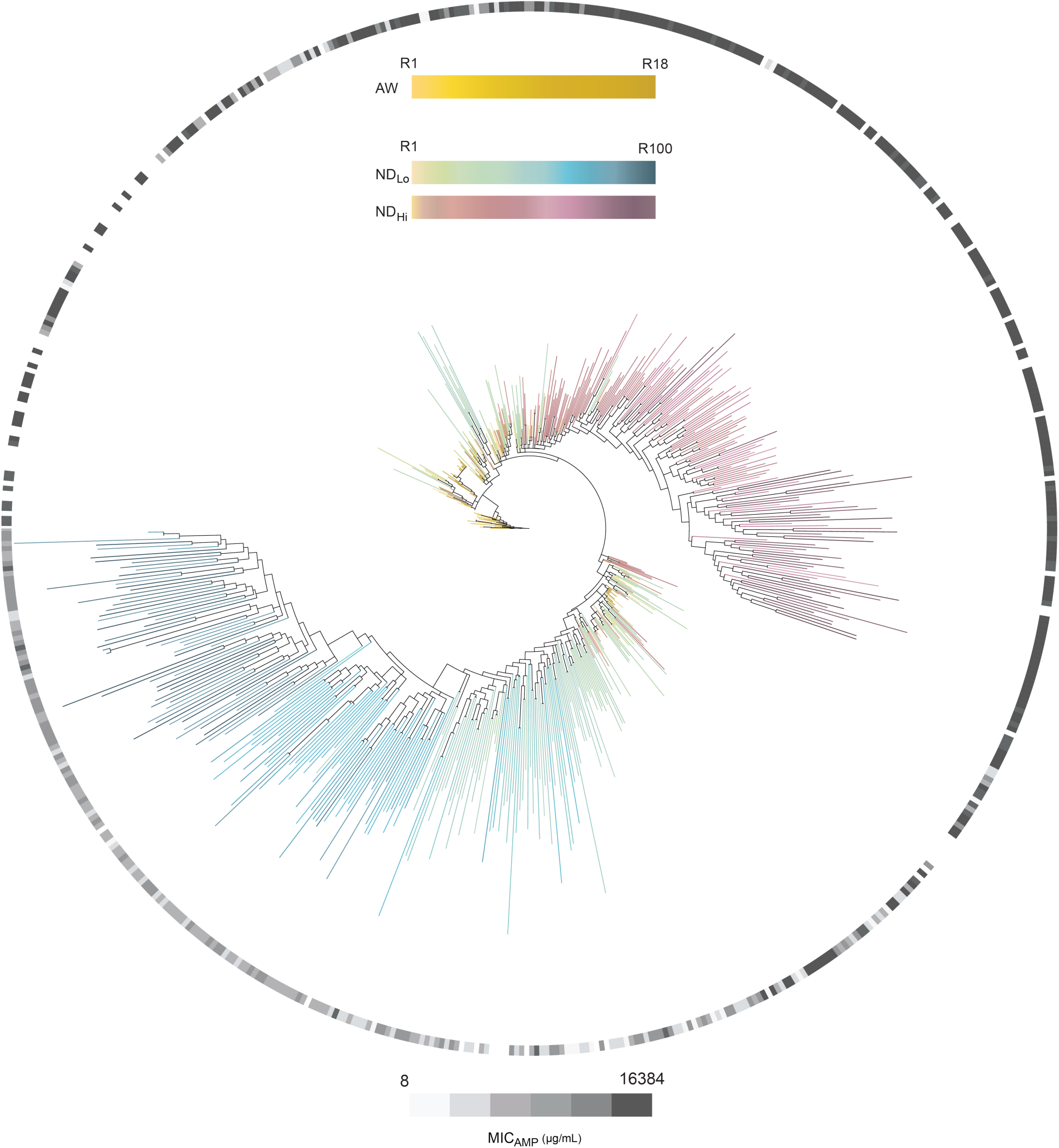
Phylogenetic tree of AW, NDLo and NDHi variants and their ampicillin resistance levels. Phylogenetic tree constructed from the nucleotide sequences of 24-96 individual variants randomly picked from the AW (yellow-orange) NDLo (light green-dark blue) and NDHi (orange-magenta) libraries, where the MICAMP of each variant is color-coded by shades of grey (white: no MIC data; light grey: 8 µg/mL; dark grey = 16,384 μg/mL). The sequences used for the tree can be found and their association with each ampicillin MIC can be found in **Supplementary Data 2**, and all DNA sequences can be found in **Supplementary Data 8**.

**Extended Data Fig. 7.**
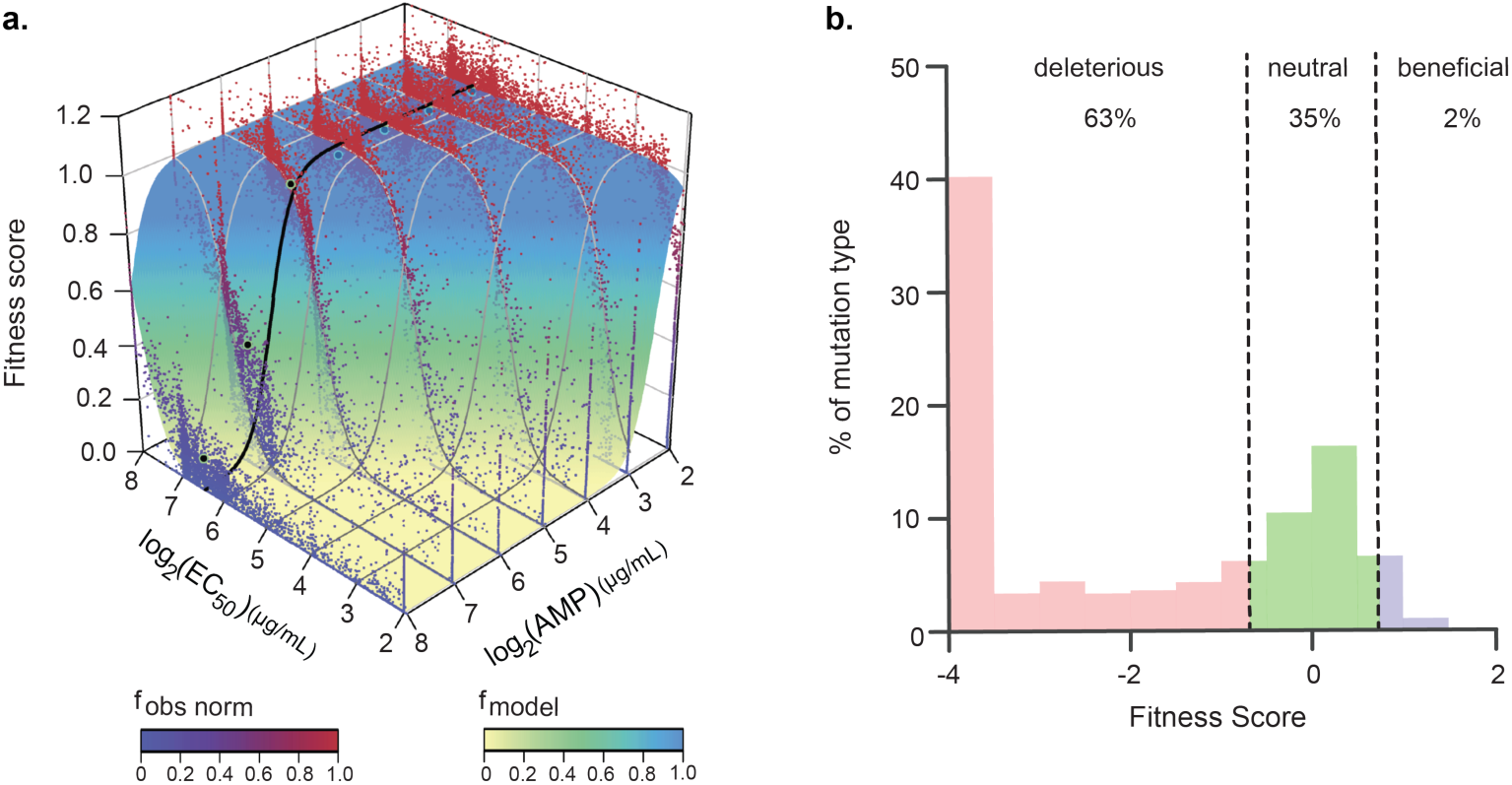
The wtVIM-2 phenotype-fitness landscape and DME. **a,** The wtVIM-2 phenotype-fitness landscape as a function of EC50 and AMP concentration is shown. The dots (‘fobs norm’) reflects the observed fitness score is plotted in relation to the AMP concentration during selection, and the surface (‘fmodel’) reflects modelled fitness scores of the single point mutants across different AMP concentrations. *Adapted from Chen et al., 202*1^26^. **b,** Distribution of fitness effects for all single amino acid variants of *wt*VIM-2. The vertical grey lines indicate fitness score (f-score) cut-offs used to classify fitness effects as positive (0.7 < f-score < 4), neutral ( -0.7 ≤ f-score ≤ 0.7), or negative ( -4 < f-score < -0.7). (The fitness scores for each single VIM-2 mutant were calculated where the selective ampicillin concentration is equal to the EC50 of *wt*VIM-2, was used to calculate the effect of each mutation on EC50, shown in fig. MS1.) *Adapted from Chen, et al.,*2020^25^.

**Extended Data Fig. 8.**
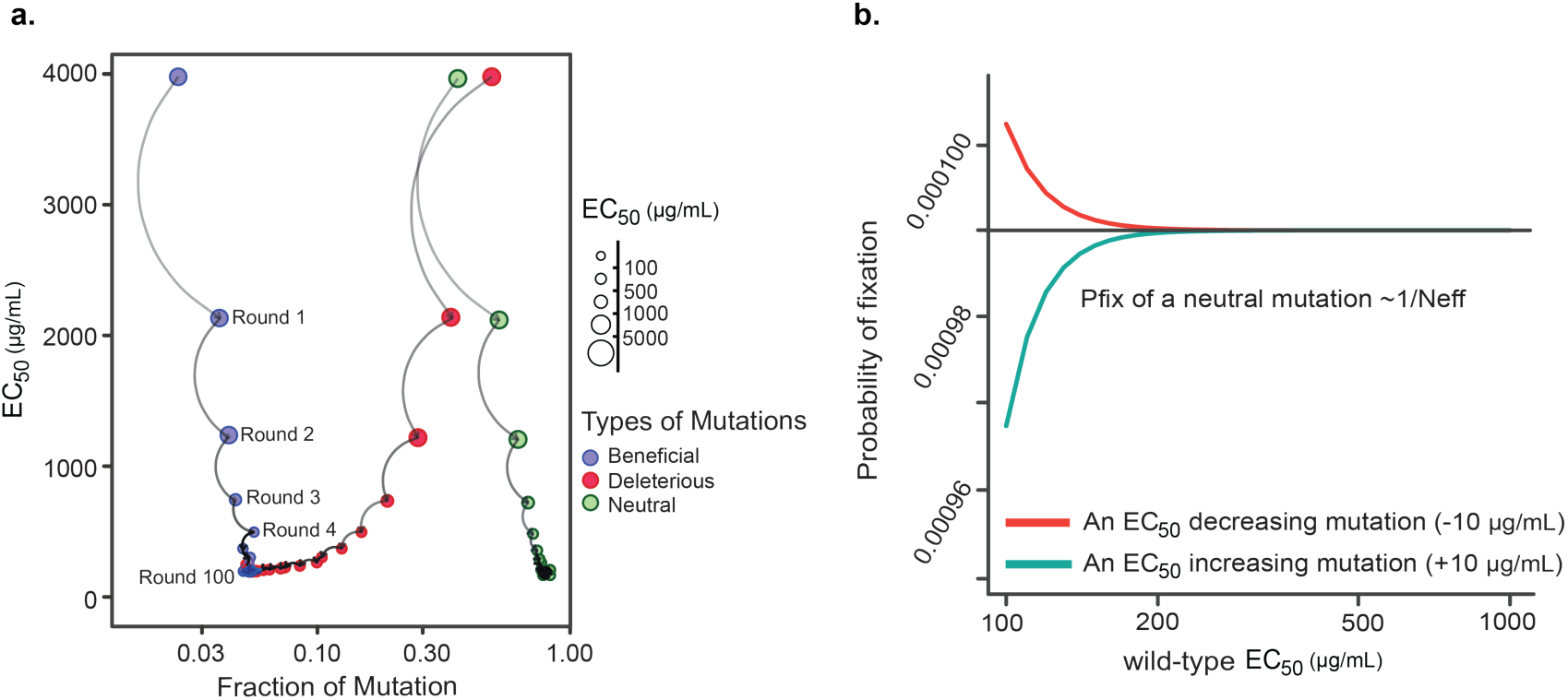
Mutation-selection balance in NDLo and probability of fixation of mutations. **a,** Established mutation-selection balance in the antibiotic-resistance level of the simulated NDLo trajectory population. The fraction of fixated mutations that are beneficial (blue), deleterious (red) and neutral (green) on the fitness of evolved enzyme variants from the NDLo trajectory as a function of the average ampicillin EC50 of each library. The size of the dots reflects the average EC50 of the library. The product of the effective population size and the selection coefficient (NeffX*s*) is used to count the fraction of deleterious (NeffX*s* < -1), beneficial (NeffX*s* > 1), and neutral (|NeffX*s*|*<* 1) substitutions. For all simulations, Neff is 10^4^ which is ∼ equal to the number of colonies sampled from LB-agar plates in each generation. Data and calculations used for the plot can be found in **Supplementary Data 6** (effects of mutations on fitness and EC50) and **7** (percentages of mutations categorized based on their effect on EC50 and fitness). **b,** Probability of fixation of mutations in the NDLo trajectory as a function of variant ampicillin EC50. The probability of fixation of a mutation that either increases (red) or decreases (blue) the ampicillin EC50 of a variant evolving under the NDLo trajectory is plotted for variants with different EC50 values. The probability of fixation of mutations that decrease or increase ampicillin EC50 of a variant with an EC50 above ∼200 ug/mL becomes indistinguishable from neutral mutations (Pfix ∼ 1/Neff).

**Extended Data Fig. 9.**
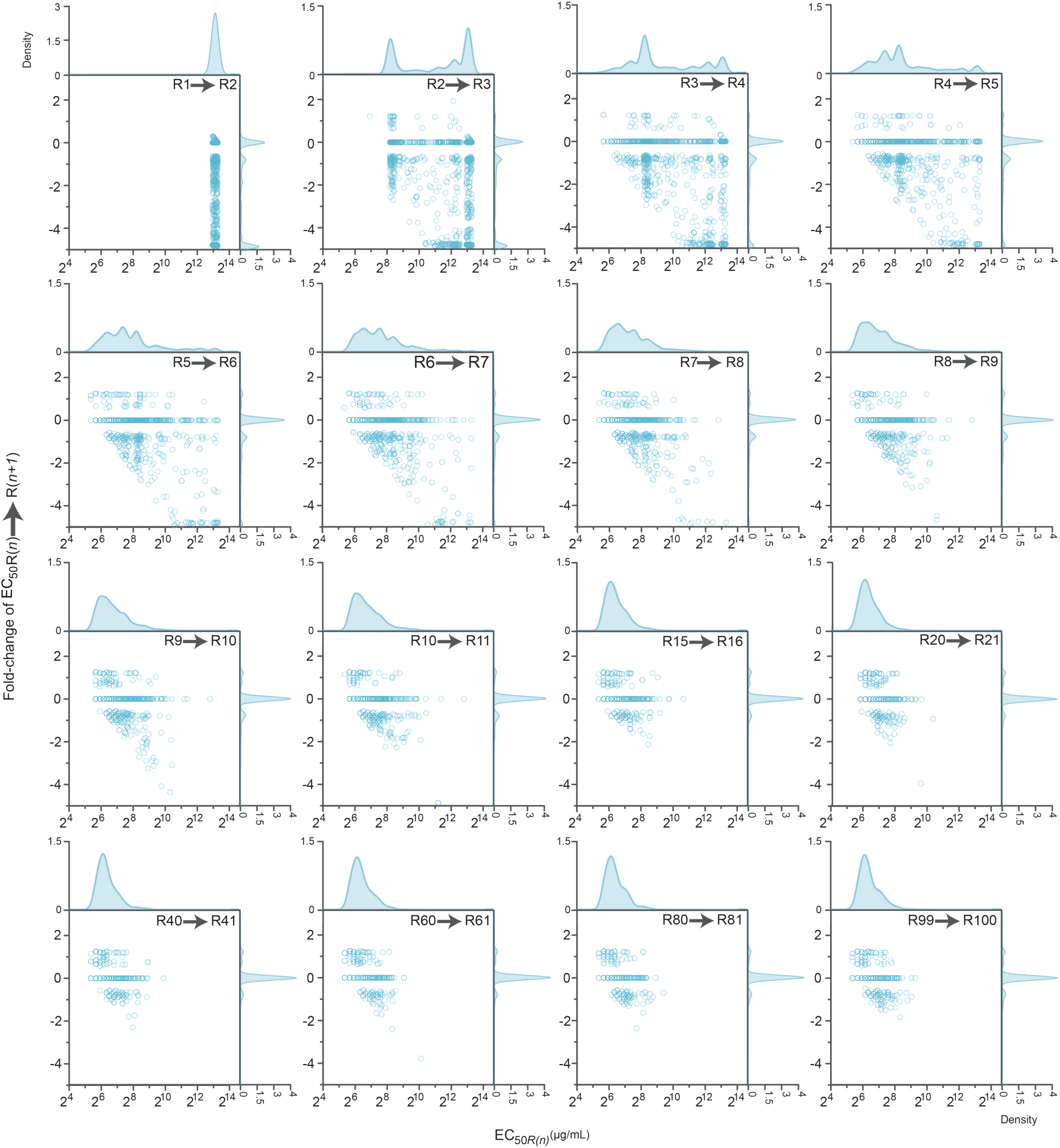
DME on the ampicillin EC50 of VIM-2 variants in the simulation of the NDLo evolution trajectory. Scatter plots show the distribution of mutational effects on the ampicillin EC50 of variants evolving under the simulated NDLo regime, in comparison to the ampicillin EC50 of the 10^3^ variants before acquiring the mutation. The mutational effects on resistance level are shown for the simulated variants as they move from R(*n*) to R(*n+1*). The change was calculated as the log2-scale fold-change in the EC50 of each variant at round (*n+*1) compared to the previous round (*n*). The gaussian kernel density distribution of the effect of mutations on resistance (*ie.,* log2 fold-change in EC50) and distribution of EC50 values of variants before acquiring the mutation is shown on the top and right axes in lighter grey (bandwidth=0.02). Data and calculations can be found in **Supplementary Data 6**. The relative percentage of each type of mutation based on its effect on fitness and EC50 can be found in **Supplementary Data 7.**

**Extended Data Fig. 10.**
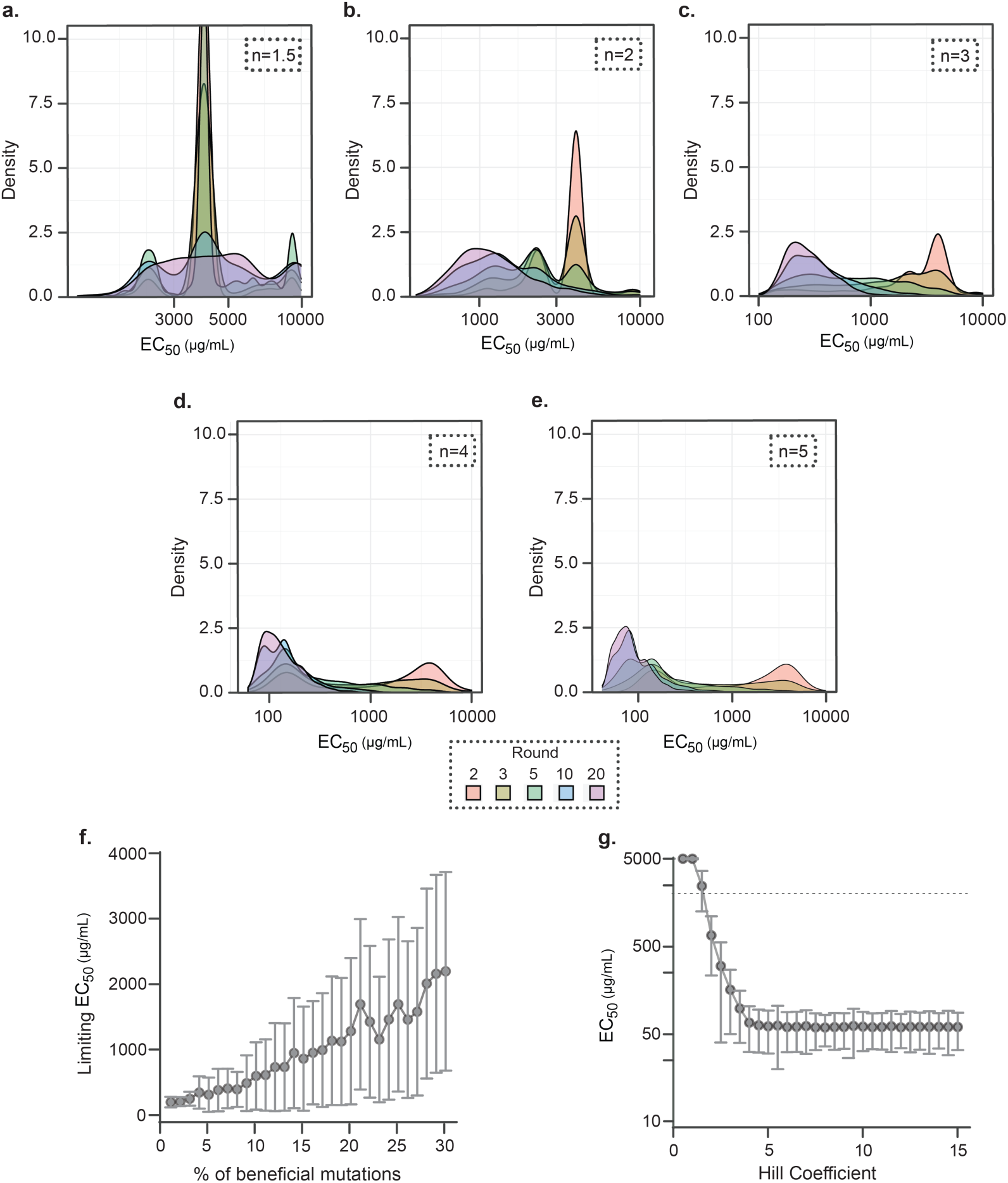
Simulation of the NDLo evolution trajectory under different conditions and sensitivity of equilibrium values to parameters. **a-e,** Simulated distribution of EC50 values of variants evolving under the NDLo trajectory with different hill coefficients: n =1.5 (**a**), n =2 (**b)**, n =3 (**c**), n =4 (**d**), n =5 (**e**); cross 20 rounds of mutagenesis and selection. **f, g,** Relationship between the percentage of positive mutations available and the limiting average EC50 and its variation (**f**), and the dose-response growth curve hill coefficient of individual VIM-2 variants and the average EC50 (**g**) of the population when it stabilizes at mutation-selection balance on a threshold-like landscape.

## Supplementary Tables

**Supplementary Table 1.** Resistance phenotype analysis of the AW, NDHi and NDLo libraries. The average ampicillin resistance level (EC50) and within-population phenotypic diversity (calculated by dividing the EC90 and EC10 for ampicillin) of the experimental evolution libraries, alongside their SEM values are shown derived from ampicillin dose response assays in (µg/mL). ‘n’ reflects the number of independent dose-response assays conducted for each library. For the raw data used to calculate these values, refer to **Supplementary Data 4.**

|  | n | EC<sub>50</sub> | EC<sub>10</sub> | EC<sub>90</sub> | EC<sub>90</sub> / EC<sub>10</sub> |
| --- | --- | --- | --- | --- | --- |
| wtVIM-2 | 4 | 93 ± 7.3 | 62 ± 5.9 | 140 ± 11 | 2.3 ± 0.14 |
| **AW** |  |  |  |  |  |
| Round | n | EC<sub>50</sub> | EC<sub>10</sub> | EC<sub>90</sub> | EC<sub>90</sub> / EC<sub>10</sub> |
| 1 |  | 270 ± 3 | 160 ± 7 | 460 ± 9 | 2.9 ± 0.17 |
| 2 |  | 780 ± 40 | 546 ± 30 | 1110 ± 50 | 2.0 ± 0.02 |
| 3 |  | 1000 ± 10 | 898 ± 20 | 1120 ± 10 | 1.3 ± 0.02 |
| 6 | 2 | 2200 ± 10 | 1753 ± 120 | 2720 ± 170 | 1.6 ± 0.20 |
| 9 |  | 2600 ± 210 | 1880 ± 30 | 3650 ± 500 | 1.9 ± 0.23 |
| 12 |  | 2600 ± 40 | 1534 ± 60 | 4270 ± 300 | 2.8 ± 0.30 |
| 15 |  | 3400 ± 230 | 2315 ± 180 | 4980 ± 290 | 2.2 ± 0.04 |
| 18 |  | 3900 ± 290 | 2758 ± 280 | 5540 ± 250 | 2.0 ± 0.12 |
| **ND<sub>Lo</sub>** |  |  |  |  |  |
| Round | n | EC<sub>50</sub> | EC<sub>10</sub> | EC<sub>90</sub> | EC<sub>90</sub> / EC<sub>10</sub> |
| 1 |  | 2310 ± 260 | 1110 ± 140 | 4900 ± 200 | 4.7 ± 0.75 |
| 2 |  | 1500 ± 140 | 390 ± 20 | 5970 ± 840 | 15 ± 1.7 |
| 5 |  | 410 ± 140 | 22 ± 4.9 | 8780 ± 2010 | 550 ± 260 |
| 10 |  | 160 ± 40 | 12 ± 3.3 | 2120 ± 191 | 200 ± 40 |
| 20 |  | 56 ± 23 | 9 ± 2.1 | 380 ± 24 | 51 ± 9.9 |
| 30 |  | 84 ± 7.3 | 12 ± 1.4 | 500 ± 70 | 52 ± 2.9 |
| 40 | 4 | 51 ± 9.6 | 11 ± 1.3 | 250 ± 24 | 24 ± 0.6 |
| 50 |  | 39 ± 5.6 | 7 ± 1.2 | 220 ± 42 | 32 ± 1.9 |
| 60 |  | 55 ± 7.1 | 17 ± 2.3 | 180 ± 20 | 11 ± 0.5 |
| 70 |  | 54 ± 6.7 | 19 ± 2.1 | 160 ± 16 | 8.4 ± 0.25 |
| 80 |  | 55 ± 5.7 | 17 ± 1.2 | 180 ± 15 | 11 ± 0.5 |
| 90 |  | 66 ± 4.0 | 18 ± 1.4 | 250 ± 32 | 14 ± 2.0 |
| 100 |  | 81 ± 4.8 | 22 ± 1.5 | 300 ± 58 | 13 ± 2.0 |
| **ND<sub>Hi</sub>** |  |  |  |  |  |
| Round | n | EC<sub>50</sub> | EC<sub>10</sub> | EC<sub>90</sub> | EC<sub>90</sub> / EC<sub>10</sub> |
| 1 |  | 2760 ± 10 | 1800 ± 200 | 4130 ± 350 | 2.3 ± 0.04 |
| 2 |  | 2990 ± 260 | 2020 ± 390 | 4430 ± 760 | 2.2 ± 0.05 |
| 5 |  | 2700 ± 550 | 1730 ± 250 | 4220 ± 510 | 2.5 ± 0.05 |
| 10 |  | 2190 ± 360 | 1310 ± 280 | 3690 ± 350 | 2.9 ± 0.36 |
| 20 |  | 2110 ± 340 | 1210 ± 290 | 3700 ± 240 | 3.2 ± 0.58 |
| 30 |  | 2300 ± 330 | 1250 ± 430 | 4260 ± 880 | 3.6 ± 0.54 |
| 40 | 2 | 2380 ± 630 | 1300 ± 390 | 4350 ± 960 | 3.4 ± 0.29 |
| 50 |  | 1770 ± 620 | 910 ± 300 | 3440 ± 890 | 3.9 ± 0.28 |
| 60 |  | 2420 ± 520 | 1350 ± 520 | 4360 ± 1200 | 3.4 ± 0.44 |
| 70 |  | 2490 ± 790 | 1440 ± 440 | 4300 ± 910 | 3.1 ± 0.30 |
| 80 |  | 2800 ± 640 | 1780 ± 460 | 4500 ± 1100 | 2.5 ± 0.07 |
| 90 |  | 2960 ± 700 | 1900 ± 200 | 4600 ± 1100 | 2.4 ± 0.30 |
| 100 |  | 2690 ± 500 | 1590 ± 460 | 4580 ± 740 | 3.0 ± 0.40 |

**Supplementary Table 2.** Phenotypic analysis of individual variants from the AW, NDHi and NDLo libraries. The averages ampicillin minimum inhibitory concentration and the coefficient of variation of MIC values of experimental evolution libraries are shown in (µg/mL). The coefficient of variation of MIC values of individual variants from the same library is used to estimate within-library phenotypic diversity. For the raw data used to calculate these values, refer to **Supplementary Data 3.**

|  | n | Mean |  | CV |
| --- | --- | --- | --- | --- |
| *wt*VIM-2 | 2 | 100 | ± 10 | 0.5 |
| AW |  |  |  |  |
| Round | n | Mean |  | CV |
| 3 | 10 | 4900 | ± 600 | 0.5 |
| 6 | 10 | 8200 | ± 0 | 0.0 |
| 9 | 11 | 14000 | ± 1000 | 0.3 |
| 18 | 29 | 11000 | ± 800 | 0.4 |
| ND<sub>Lo</sub> |  |  |  |  |
| Round | n | Mean |  | CV |
| 1 | 24 | 7300 | ± 1000 | 0.8 |
| 2 | 24 | 4500 | ± 1000 | 1.0 |
| 5 | 24 | 1400 | ± 500 | 2.0 |
| 10 | 48 | 540 | ± 100 | 2.0 |
| 15 | 48 | 460 | ± 100 | 2.0 |
| 20 | 48 | 440 | ± 100 | 2.0 |
| 30 | 23 | 600 | ± 100 | 1.0 |
| 40 | 72 | 250 | ± 40 | 1.0 |
| 50 | 48 | 160 | ± 50 | 2.0 |
| 60 | 92 | 270 | ± 30 | 1.0 |
| 70 | 47 | 390 | ± 40 | 0.6 |
| 80 | 71 | 360 | ± 50 | 1.0 |
| 90 | 48 | 390 | ± 40 | 0.8 |
| 100 | 168 | 780 | ± 50 | 0.9 |
| ND<sub>Hi</sub> |  |  |  |  |
| Round | n | Mean |  | CV |
| 1 | 24 | 9900 | ± 1000 | 0.8 |
| 2 | 24 | 9600 | ± 1000 | 1.0 |
| 5 | 24 | 9900 | ± 500 | 2.0 |
| 10 | 48 | 8000 | ± 100 | 2.0 |
| 15 | 48 | 7300 | ± 100 | 2.0 |
| 20 | 48 | 9100 | ± 100 | 2.0 |
| 30 | 23 | 12000 | ± 100 | 1.0 |
| 40 | 72 | 7900 | ± 40 | 1.0 |
| 50 | 48 | 6200 | ± 50 | 2.0 |
| 60 | 92 | 8700 | ± 30 | 1.0 |
| 70 | 47 | 8400 | ± 40 | 0.6 |
| 80 | 71 | 8100 | ± 50 | 1.0 |
| 90 | 48 | 9300 | ± 40 | 0.8 |
| 100 | 93 | 14000 | ± 50 | 0.9 |

**Supplementary Table 3.** Genotypic analysis of the AW, NDHi and NDLo libraries. Mean and standard error of nucleotide mutations (transitions and transversions) and amino acid mutations in the signal peptide and the mature protein; Na/(Nt), percent sequence identity shared with *wt*VIM-2 and within the library are given for 24-96 individual variants from the evolution libraries. For the raw data used to calculate these values, refer to **Supplementary Data 2.**

| AW |  |  |  |  |  |  |  |  |  |  |  |  |  |  |  |  |  |
| --- | --- | --- | --- | --- | --- | --- | --- | --- | --- | --- | --- | --- | --- | --- | --- | --- | --- |
| Nucleotide Mutations |  |  |  |  |  |  |  | Amino Acid Mutations |  |  |  | Na/Nt |  | Percent Identity to wtVIM-2 |  | Percent Identity within Library |  |
| Round | n | Transition |  | Transversion |  | Total |  | Signal Peptide |  | Protein |  |  |  |  |  |  |  |
|  |  | Mean | SEM | Mean | SEM | Mean | SEM | Mean | SEM | Mean | SEM | Mean | SEM | Mean | SEM | Mean | SEM |
| 3 | 19 | 9 | ± 0.6 | 6 | ± 0.5 | 15 | ± 0.8 | 3 | ± 0.2 | 8 | ± 0.3 | 0.52 | ± 0.03 | 97.1 | ± 0.1 | 95.8 | ± 0.1 |
| 6 | 17 | 12 | ± 0.6 | 7 | ± 0.7 | 19 | ± 0.8 | 3 | ± 0.2 | 10 | ± 0.4 | 0.53 | ± 0.02 | 96.2 | ± 0.2 | 96.5 | ± 0.1 |
| 9 | 21 | 19 | ± 0.5 | 9 | ± 0.1 | 28 | ± 0.5 | 3 | ± 0.1 | 14 | ± 0.2 | 0.48 | ± 0.01 | 94.9 | ± 0.1 | 99.6 | ± 0.0 |
| 12 | 17 | 23 | ± 0.5 | 10 | ± 0.4 | 32 | ± 0.1 | 4 | ± 0.1 | 16 | ± 0.5 | 0.49 | ± 0.01 | 94.0 | ± 0.2 | 97.1 | ± 0.1 |
| 15 | 18 | 27 | ± 0.5 | 10 | ± 0.4 | 36 | ± 0.8 | 5 | ± 0.2 | 19 | ± 0.4 | 0.51 | ± 0.01 | 93.1 | ± 0.2 | 97.0 | ± 0.1 |
| 18 | 23 | 31 | ± 0.6 | 12 | ± 0.6 | 43 | ± 0.8 | 5 | ± 0.1 | 21 | ± 0.5 | 0.50 | ± 0.01 | 91.9 | ± 0.2 | 95.2 | ± 0.1 |
| ND<sub>Lo</sub> |  |  |  |  |  |  |  |  |  |  |  |  |  |  |  |  |  |
| Nucleotide Mutations |  |  |  |  |  |  |  | Amino Acid Mutations |  |  |  | Na/Nt |  | Percent Identity to wtVIM-2 |  | Percent Identity within Library |  |
| Round | n | Transition |  | Transversion |  | Total |  | Signal Peptide |  | Protein |  |  |  |  |  |  |  |
|  |  | Mean | SEM | Mean | SEM | Mean | SEM | Mean | SEM | Mean | SEM | Mean | SEM | Mean | SEM | Mean | SEM |
| 1 | 24 | 34 | ± 0.7 | 13 | ± 0.6 | 47 | ± 0.9 | 6 | ± 0.2 | 22 | ± 0.5 | 0.48 | ± 0.01 | 91.6 | ± 0.2 | 94.2 | ± 0.1 |
| 2 | 24 | 33 | ± 0.8 | 13 | ± 0.5 | 46 | ± 0.9 | 6 | ± 0.2 | 22 | ± 0.5 | 0.49 | ± 0.01 | 91.6 | ± 0.2 | 94.0 | ± 0.1 |
| 5 | 21 | 35 | ± 0.9 | 16 | ± 0.8 | 51 | ± 1.2 | 6 | ± 0.3 | 26 | ± 0.7 | 0.51 | ± 0.01 | 90.3 | ± 0.2 | 91.5 | ± 0.1 |
| 10 | 21 | 45 | ± 1.0 | 18 | ± 0.8 | 63 | ± 1.3 | 8 | ± 0.4 | 31 | ± 0.7 | 0.50 | ± 0.01 | 88.2 | ± 0.3 | 88.4 | ± 0.1 |
| 15 | 23 | 49 | ± 1.0 | 20 | ± 0.7 | 69 | ± 1.1 | 9 | ± 0.3 | 33 | ± 0.7 | 0.48 | ± 0.01 | 87.6 | ± 0.3 | 87.5 | ± 0.1 |
| 20 | 22 | 54 | ± 1.2 | 23 | ± 0.7 | 76 | ± 1.2 | 9 | ± 0.2 | 37 | ± 0.9 | 0.49 | ± 0.01 | 86.0 | ± 0.3 | 85.9 | ± 0.1 |
| 30 | 25 | 60 | ± 1.2 | 27 | ± 0.8 | 87 | ± 1.3 | 10 | ± 0.3 | 41 | ± 0.9 | 0.47 | ± 0.01 | 84.6 | ± 0.3 | 83.6 | ± 0.1 |
| 40 | 29 | 69 | ± 0.9 | 28 | ± 0.8 | 97 | ± 1.0 | 11 | ± 0.4 | 46 | ± 0.7 | 0.48 | ± 0.01 | 82.6 | ± 0.2 | 80.7 | ± 0.1 |
| 50 | 18 | 79 | ± 1.1 | 35 | ± 1.0 | 114 | ± 1.7 | 13 | ± 0.6 | 52 | ± 1.2 | 0.46 | ± 0.01 | 80.3 | ± 0.4 | 78.4 | ± 0.2 |
| 60 | 37 | 83 | ± 1.1 | 36 | ± 0.8 | 118 | ± 1.2 | 11 | ± 0.3 | 51 | ± 0.7 | 0.43 | ± 0.00 | 80.6 | ± 0.3 | 80.9 | ± 0.1 |
| 70 | 22 | 90 | ± 1.5 | 37 | ± 2.0 | 128 | ± 1.6 | 12 | ± 0.5 | 54 | ± 0.8 | 0.42 | ± 0.00 | 79.6 | ± 0.3 | 80.6 | ± 0.2 |
| 80 | 23 | 65 | ± 1.6 | 40 | ± 1.1 | 135 | ± 1.7 | 12 | ± 0.3 | 60 | ± 0.9 | 0.44 | ± 0.00 | 77.4 | ± 0.3 | 80.4 | ± 0.2 |
| 90 | 21 | 97 | ± 1.2 | 45 | ± 1.2 | 142 | ± 1.6 | 14 | ± 0.4 | 60 | ± 1.1 | 0.42 | ± 0.01 | 76.9 | ± 0.4 | 80.5 | ± 0.2 |
| 100 | 23 | 99 | ± 1.7 | 46 | ± 1.0 | 145 | ± 1.8 | 14 | ± 0.4 | 61 | ± 1.2 | 0.42 | ± 0.01 | 76.4 | ± 0.4 | 79.5 | ± 0.1 |
| ND<sub>Hi</sub> |  |  |  |  |  |  |  |  |  |  |  |  |  |  |  |  |  |
| Nucleotide Mutations |  |  |  |  |  |  |  | Amino Acid Mutations |  |  |  | Na/Nt |  | Percent Identity to wtVIM-2 |  | Percent Identity within Library |  |
| Round | n | Transition |  | Transversion |  | Total |  | Signal Peptide |  | Protein |  |  |  |  |  |  |  |
|  |  | Mean | SEM | Mean | SEM | Mean | SEM | Mean | SEM | Mean | SEM | Mean | SEM | Mean | SEM | Mean | SEM |
| 5 | 22 | 35 | ± 0.8 | 14 | ± 0.6 | 49 | ± 0.7 | 6 | ± 0.3 | 24 | ± 0.5 | 0.49 | ± 0.01 | 91.5 | ± 0.4 | 94.3 | ± 0.1 |
| 10 | 34 | 42 | ± 0.9 | 16 | ± 0.5 | 59 | ± 0.9 | 7 | ± 0.2 | 28 | ± 0.6 | 0.47 | ± 0.01 | 89.8 | ± 0.3 | 92.1 | ± 0.1 |
| 15 | 21 | 46 | ± 0.4 | 19 | ± 0.7 | 65 | ± 1.6 | 7 | ± 0.2 | 30 | ± 0.7 | 0.46 | ± 0.01 | 88.7 | ± 0.3 | 91.0 | ± 0.1 |
| 20 | 28 | 51 | ± 1.0 | 20 | ± 0.7 | 71 | ± 1.1 | 8 | ± 0.3 | 31 | ± 0.8 | 0.44 | ± 0.01 | 88.1 | ± 0.3 | 90.8 | ± 0.1 |
| 30 | 30 | 59 | ± 1.0 | 25 | ± 0.7 | 84 | ± 0.9 | 9 | ± 0.4 | 35 | ± 0.5 | 0.42 | ± 0.01 | 86.8 | ± 0.2 | 88.7 | ± 0.1 |
| 40 | 31 | 63 | ± 0.9 | 25 | ± 0.7 | 88 | ± 1.2 | 10 | ± 0.4 | 37 | ± 0.7 | 0.42 | ± 0.01 | 86.0 | ± 0.3 | 88.2 | ± 0.1 |
| 50 | 20 | 73 | ± 1.4 | 31 | ± 1.0 | 103 | ± 1.3 | 9 | ± 0.5 | 41 | ± 0.9 | 0.40 | ± 0.01 | 84.8 | ± 0.4 | 86.3 | ± 0.1 |
| 60 | 24 | 76 | ± 1.2 | 31 | ± 1.0 | 107 | ± 1.4 | 11 | ± 0.3 | 43 | ± 0.7 | 0.41 | ± 0.01 | 83.8 | ± 0.2 | 85.9 | ± 0.1 |
| 80 | 28 | 86 | ± 1.3 | 38 | ± 1.2 | 124 | ± 1.5 | 12 | ± 0.4 | 47 | ± 0.8 | 0.38 | ± 0.00 | 81.4 | ± 0.4 | 83.3 | ± 0.1 |
| 100 | 20 | 98 | ± 1.1 | 43 | ± 1.3 | 141 | ± 2.0 | 13 | ± 0.6 | 52 | ± 1.1 | 0.37 | ± 0.01 | 80.4 | ± 0.4 | 82.7 | ± 0.2 |

**Supplementary Table 4.** Population-level phenotypic analysis of the NDLo-wt libraries. The median ampicillin resistance level (EC50), and within-library phenotypic diversity (calculated by dividing the EC90 and EC10 for ampicillin of the experimental evolution libraries, alongside their SEM values are shown derived from ampicillin dose response assays in (µg/mL). ‘n’ reflects the number of independent dose-response assays conducted for each library. For the raw data used to calculate these values, refer to **Supplementary Data 4.**

|  | n | EC<sub>50</sub> | EC<sub>10</sub> | EC<sub>90</sub> | EC<sub>90</sub> / EC<sub>10</sub> |
| --- | --- | --- | --- | --- | --- |
| wtVIM-2 | 2 | 134 ± 9 | 103 ± 5 | 177 ± 37 | 1.7 ± 0.45 |
| **ND<sub>Lo-wt</sub>VIM-2** |  |  |  |  |  |
| Round | n | EC<sub>50</sub> | EC<sub>10</sub> | EC<sub>90</sub> | EC<sub>90</sub> / EC<sub>10</sub> |
| 1 | 2 | 110 ± 3 | 56 ± 12 | 225 ± 38 | 4.3 ± 1.6 |
| 2 | 3 | 95 ± 5 | 48 ± 5 | 191 ± 21 | 4.1 ± 0.8 |
| 3 | 2 | 88 ± 4 | 36 ± 8 | 221 ± 28 | 6.6 ± 2.3 |
| 4 | 3 | 79 ± 4 | 31 ± 2 | 206 ± 18 | 6.8 ± 0.8 |
| 5 | 2 | 79 ± 4 | 27 ± 2 | 232 ± 3 | 8.6 ± 0.5 |
| 6 | 3 | 65 ± 7 | 19 ± 3 | 221 ± 10 | 12 ± 2.1 |
| 7 | 1 | 75 | 25 | 226 | 9.1 |
| 8 | 2 | 73 ± 13 | 24 ± 9 | 230 ± 8 | 11 ± 3.8 |
| 9 | 2 | 70 ± 15 | 23 ± 12 | 221 ± 1 | 13 ± 6.3 |
| 10 | 3 | 63 ± 9 | 19 ± 5 | 212 ± 11 | 12 ± 2.4 |
| 11 | 2 | 51 ± 2 | 13 | 197 ± 8 | 15 |
| 12 | 2 | 50 ± 2 | 13 ± 1 | 191 ± 5 | 15 ± 0.6 |
| 13 | 2 | 52 ± 1 | 14 | 195 ± 3 | 14 ± 0.1 |
| 14 | 2 | 60 ± 2 | 17 | 209 ± 14 | 12 ± 0.7 |
| 15 | 1 | 62 | 14 | 210 | 15 |

**Supplementary Table 5.** Genotypic analysis of the NDLo-wt libraries. Mean and standard error of nucleotide mutations (transitions and transversions) and amino acid mutations in the signal peptide and the mature protein; Na/Nt, percent sequence identity shared with *wt*VIM-2 and within the library are given for 2-15 individual variants from the evolution libraries. For the raw data used to calculate these values, refer to **Supplementary Data 2.**

| NDLo-wt |  |  |  |  |  |  |  |  |  |  |  |  |  |  |  |  |  |
| --- | --- | --- | --- | --- | --- | --- | --- | --- | --- | --- | --- | --- | --- | --- | --- | --- | --- |
| Nucleotide Mutations |  |  |  |  |  |  |  | Amino Acid Mutations |  |  |  | Na/Nt |  | Percent Identity to wtVIM-2 |  | Percent Identity within Library |  |
| Round | n | Transition |  | Transversion |  | Total |  | Signal Peptide |  | Protein |  |  |  |  |  |  |  |
|  |  | Mean | SEM | Mean | SEM | Mean | SEM | Mean | SEM | Mean | SEM | Mean | SEM | Mean | SEM | Mean | SEM |
| 1 | 5 | 1 | ± 0.3 | 1 | ± 0.4 | 2 | ± 0.6 | 0 | ± 0.0 | 1 | ± 0.7 | 0.70 | ± 0.20 | 99.5 | ± 0.3 | 98.9 | ± 0.2 |
| 2 | 3 | 3 | ± 0.9 | 2 | ± 0.7 | 4 | ± 1.2 | 0 | ± 0.0 | 3 | ± 0.9 | 0.59 | ± 0.05 | 99.0 | ± 0.3 | 98.2 | ± 0.1 |
| 3 | 5 | 3 | ± 0.8 | 1 | ± 0.9 | 4 | ± 1.3 | 0 | ± 0.4 | 3 | ± 0.9 | 0.77 | ± 0.12 | 98.9 | ± 0.3 | 97.9 | ± 0.3 |
| 4 | 6 | 5 | ± 0.8 | 2 | ± 0.6 | 7 | ± 1.2 | 1 | ± 0.3 | 4 | ± 1.1 | 0.54 | ± 0.10 | 98.5 | ± 0.4 | 97.0 | ± 0.3 |
| 5 | 9 | 5 | ± 0.7 | 1 | ± 0.3 | 7 | ± 0.6 | 1 | ± 0.4 | 4 | ± 0.5 | 0.55 | ± 0.05 | 98.6 | ± 0.2 | 97.3 | ± 0.1 |
| 6 | 15 | 6 | ± 0.8 | 3 | ± 0.5 | 9 | ± 1.0 | 2 | ± 0.3 | 6 | ± 0.7 | 0.64 | ± 0.04 | 97.8 | ± 0.3 | 95.9 | ± 0.1 |
| 7 | 14 | 8 | ± 1.3 | 3 | ± 0.4 | 11 | ± 1.5 | 2 | ± 0.4 | 6 | ± 0.9 | 0.55 | ± 0.05 | 97.8 | ± 0.3 | 96.0 | ± 0.2 |
| 8 | 2 | 7 | ± 1.0 | 5 | ± 2.0 | 12 | ± 1.0 | 1 | ± 0.5 | 9 | ± 0.5 | 0.71 | ± 0.02 | 96.8 | ± 0.2 | 93.6 | ± - |
| 10 | 14 | 10 | ± 1.1 | 5 | ± 0.8 | 15 | ± 1.4 | 2 | ± 0.4 | 8 | ± 0.9 | 0.55 | ± 0.03 | 97.0 | ± 0.3 | 93.2 | ± 1.0 |
| 12 | 3 | 16 | ± 1.0 | 6 | ± 0.3 | 22 | ± 1.3 | 5 | ± 0.7 | 11 | ± 0.0 | 0.51 | ± 0.03 | 95.9 | ± 0.0 | 92.1 | ± 0.2 |
| 15 | 5 | 19 | ± 1.9 | 6 | ± 1.4 | 24 | ± 1.7 | 4 | ± 0.9 | 14 | ± 1.5 | 0.57 | ± 0.05 | 94.5 | ± 0.6 | 91.0 | ± 0.8 |

**Supplementary Table 6.**
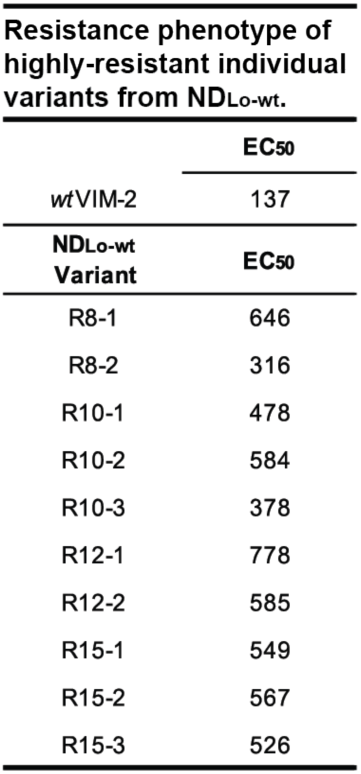
Phenotypic analysis of individual NDLo-wt trajectory variants. Ampicillin EC50 (µg/mL) values of randomly selected individual variants from R8, R10, R12 and R15 libraries of the NDL-wt trajectory that confer ampicillin resistance levels >30 higher than the selection threshold (10 µg/mL), determined by dose-response assays (Methods). The median ampicillin resistance level (EC50) of randomly selected individual variants from R8, R10, R12 and R15 libraries of the NDL-wt trajectory that confer ampicillin resistance levels >30 higher than the selection threshold (10 µg/mL) are shown in (µg/mL). For the raw data used to calculate these values, refer to **Supplementary Data 4.**

## Notes

### Competing Interest Statement

The authors have declared no competing interest.

